# Spatial continuity of neurons explains non-random network architecture

**DOI:** 10.1101/2025.08.21.671478

**Authors:** Michael W. Reimann, Daniela Egas Santander, Lida Kanari, Natalí Barros-Zulaica

**Affiliations:** Open Brain Institute, Lausanne, Switzerland; Max Planck Institute of Molecular Cell Biology and Genetics; Department of Mathematics, University of Oxford, Oxford,UK

## Abstract

Neuronal networks are characterized by complex and functionally relevant connectivity motifs. We developed an intuitive explanation for its emergence: While a class of neurons on average innervates its entire surroundings, each individual one can only cover a small part of the space. That region is different for each neuron, but not completely random, as it is physically constrained by the spatial continuity of the axon. This hypothesis was successfully tested against a morphologically-detailed model and an electron-microscopic reconstruction of cortical connectivity. We distilled it into a stochastic algorithm that generates networks, which accurately match the reference data. Our work bridges previous efforts to capture network complexity with top-down or bottom-up methods, that is, by adding complexity constraints to simple stochastic models or by predicting synapses from neuron appositions. It may improve the understanding of the impact of neuron malformations and the functional role of non-random network structure in simplified models.

## 1 Introduction

The understanding of an aspect of neural function is often derived from analyses of underlying mechanisms using experimental data, followed by extracting fundamental principles, leading to a simplified description. For the structure of neuronal connectivity at cellular resolution, i.e., the network formed by individual neurons and chemical synapses between them, this has been difficult to obtain, as most of the available data is highly subsampled. While insights into the developmental mechanisms shaping connectivity exists [1; 2] their descriptions are complex and have rarely been turned into models [3; 4; 5]. Instead, many models of circuit structure are based around high-level experimental measurements that lead to constraints on, e.g., sparsity (or connection probability) and distance-dependence [6; 7; 8]. However, these measurements come with substantial uncertainties and do not always agree with one another. For example, in a study exhaustively sampling synaptic pathways in adult mouse visual cortex, [9] found zero connectivity between pyramidal cells in layer 5. On the other hand, we found that in a recent electron microscopic reconstruction of the same system [10], connection probabilities of 6.5% are observed. Additionally, connection probabilities are inherently limited in their use to small volumes: The number of neurons in spherical volumes grows with the third power of its diameter. Hence, as volume size increases, the required precision of connection probability estimates grows rapidly, as even small probabilities represent many connections. However, progress has been made in estimating complex network properties from sparse and noisy data observations [11]. Moreover, there has been remarkable progress in software tools for neuron image segmentation and network reconstruction from electron microscopy data, steadily enabling increasingly accurate large-scale connectivity reconstructions [12; 13; 14; 15; 16].

The review by Hoffmann et al. [14] highlights the need for robust, biologically meaningful metrics to evaluate network reconstruction in large datasets, particularly those that go beyond pairwise neuronal connectivity. At the same time, experimental results have characterized many forms of *non-random higher order structure* in neuronal circuitry [17; 18; 19; 20; 21; 22; 23; 24; 25; 26] that many existing models do not capture. Models exist that take them into account: The model of Tesler et al. [27] enforces long-tailed degree distributions [28] as well as Billeh et al. [7] in its latest version (personal communication). Additionally, preferential attachment models [29; 30], developed to introduce node non-homogeneity in networks, can abstractly match scale-free degree distributions, and Brunel [31] found that such distributions already generate some of the non-random structure. However, Egas Santander et al. [25] found that they only explain a small fraction of it. On the other hand, small-world models [32; 33] generate non-random higher-order structure and distance-dependent geometric models capture spatial structure, but neither of them accurately reproduce the observed degree distributions of biological neural networks. This limitation arises because their underlying algorithms form connections either based on network properties such as node degree or rewiring probability or by incorporating explicit spatial or geometric constraints. In that regard, it has been stated that a combination of distance-dependence and constraints on attachment preference or clustered structure are needed to model realistic connectivity [34; 35; 36]. Fractal methods propose an alternative path by focusing on the (multi)fractal structures that arise in biological networks and and their generating rules, which allow one to match both degree distributions and clustering coefficients [37; 38; 11]. Remarkably, such methods have been used in networks inferred from rodent neuronal cultures, which were shown to have unique multifractal and assortative connectivity properties not captured by traditional models and have been useful for linking micro- and mesoscale connectivity [5]. However, these examples focus on inferring network structure top-down from statistical observations. It remains unsolved to link these observations to the structures arising from biophysical spatial constraints such as neuronal morphology, offering a more mechanistic understanding of connectivity formation.

A better understanding of the mechanisms of non-random connectivity is desirable, as degradation of local circuitry has been shown to be clinically relevant [39; 40]. Additionally, higher-order structure specifically, has been predicted to have an impact on the neuronal code [41] and be related to reliable responses at the level of local circuitry [25].

Our understanding of a phenomenon is often improved by considering or even simulating additional underlying details, characterizing its role, and turning it into simplified models. For example, Hodgkin-Huxley type models are simplified models of neuron excitability, but they consider individual classes of membrane-bound ion channels and their kinetics [42]. With respect to the structure of neuronal circuitry, one relevant detail is the shape of individual neurites, as proximity of an axon to another neurite (or a soma) is a required condition for the formation of a synaptic contact [43; 44]. This idea has been implemented at various levels of detail. Gandolfi et al. [28] consider the average shapes of classes of neurons, while Binzegger et al. [44]; Reimann et al. [45] use axo-dendritic appositions between individual morphologies as potential connections. Oberlaender et al. [46]; Udvary et al. [47] favor an intermediate approach that is also using individual morphology reconstructions, but calculate overlap at reduced resolution, i.e., based on the amount of neurite found in voxels. While it is still discussed how much neuron morphology shapes connectivity [48], it has been repeatedly demonstrated that models of connectivity based on axo-dendritic appositions recreate many relevant features of non-random higher order structure found in biological neuronal networks [49; 47].

We argue that the key to understanding the non-random structure of connectivity and accurately modeling it lies in its emergence from axo-dendritic appositions. We postulate a hypothesis about why it emerges and prove predictions arising from it in a connectome that has been built from morphological appositions and an electron microscopic reconstruction of circuitry in mouse V1 [10], while relevant control models led to negative results. This also results in a novel analysis that measures how strongly a connectome is affected by the hypothesized mechanism. Finally, we present a simplified connectome model that captures the essence of the proposed mechanism, but does not use reconstructed neuron morphologies. I n the model, both the spatial structure and non-random micro-structure emerge together from a common mechanism. While we have focused our analyses in this work on excitatory connections in rodent neocortical circuitry, we believe that it will generalize well to other organisms and brain regions, and we provide discussions of how this may be achieved in the future. The model is computationally cheap and can be easily scaled up, making it suitable for simulations of point neuron networks. Moreover, we sketched out an extension to long-range connectivity that may in the future make it possible to model whole-brain connectivity at cellular resolution.

## 2 Results

### 2.1 Central hypothesis: Neuron Physicality Shapes Network Structure

Several aspects of non-random connectivity have been characterized in biological neuronal networks. Reciprocal connections have been found to be more abundant than expected (Fig. 1A1, [17; 18]), although this has not been confirmed in human cortical circuitry [26]. Additionally, the connectivity is often highly clustered (Fig. 1A2), and more generally, specific “connectivity motifs” are overexpressed (Fig. 1A3; [17; 26]). Recently, it has also been found that the locations of reciprocal connections are non-random: They are found predominately within a class of connectivity motifs called “directed simplices”, a cluster of neurons that are all-to-all connected in feedforward fashion (Fig. 1A4, Fig. S7A; [25]).

**Figure 1:**
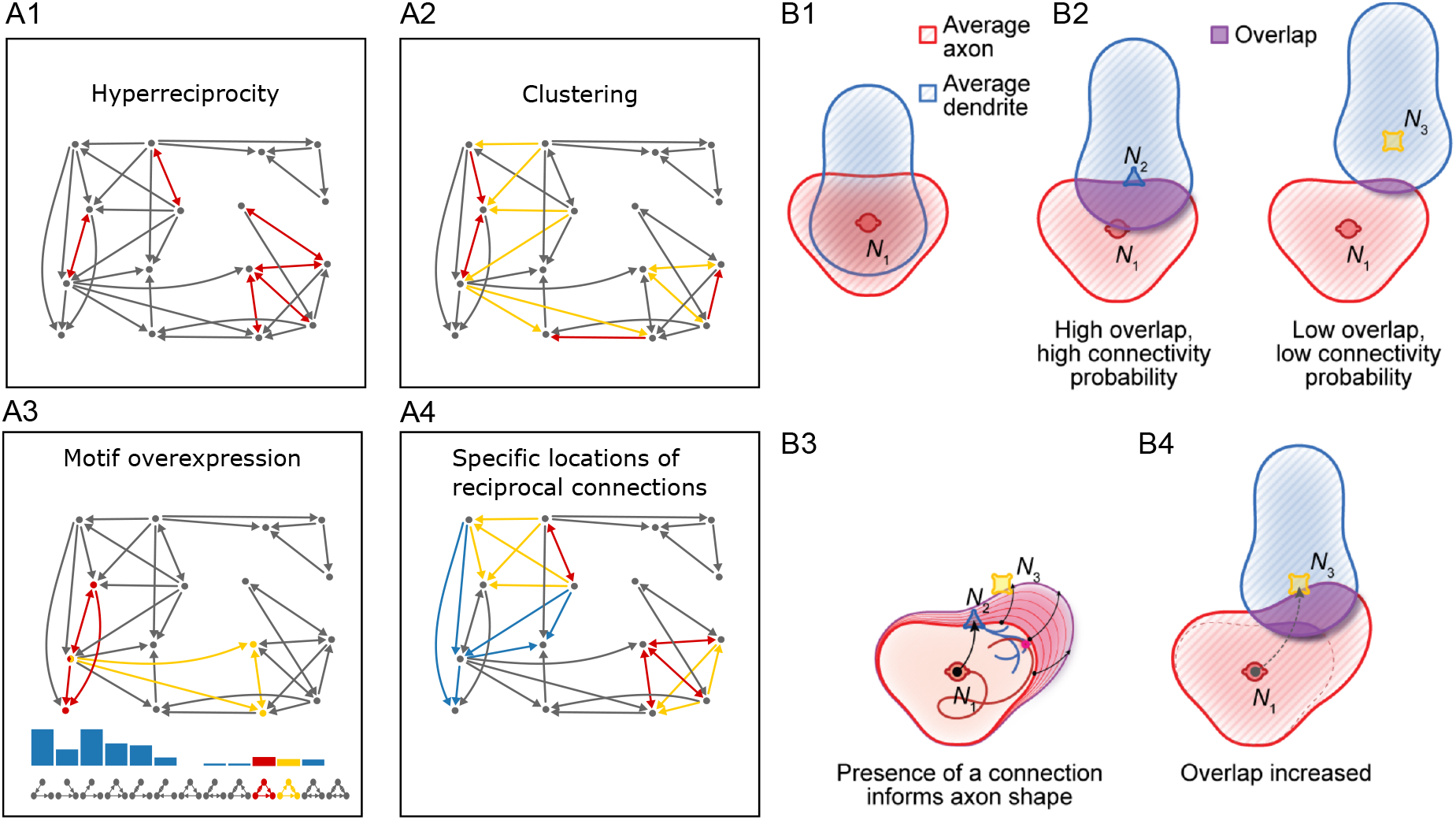
Hypothesis: Neurite physicality produce non-independent connections, leading to non-random connectivity A: Schemas of non-random trends found in neuronal connectivity illustrated on a toy network. A1: In many circuits, reciprocal connections (red) occur more often than expected by chance. A2: Connectivity is often clustered, e.g., pairs of neurons that are innervated by a common third neuron (yellow edges) tend to be connected themselves (red). A3: More general, when counting the instances of all possible connected motifs of three neurons (A3, bottom), specific motifs tend to occur more often than expected (blue bars show counts of the indicated motifs). Instances of two exemplary motifs are highlighted in red and yellow. A4: Reciprocal connections have been found to be located preferably within larger “directed simplex” motifs (see Fig S7). Two all-to-all connected cliques forming three-dimensional simplices in the toy network are indicated in yellow with reciprocal edges within them in red (top left and bottom right corner). Four out of six reciprocal connection (see A1) are within these simplices of dimension three. In blue, two-dimensional simplices with no reciprocal connections. B: Hypothesized reason: B1: For a given class of neurons, we can consider the shapes of the *average axon and dendrite*, i.e., the probability distribution for axonal and dendritic mass over all neurons in the class. B2: Depending on the offset between a pair of neurons, the average shapes can overlap strongly (left) or weakly (right), resulting in a high or low probability that axon meets dendrite, a prerequisite of a synaptic connection. This leads to a complex, offset-dependent connection probability function that is often used in models. B3: However, once a connection to a neuron is confirmed, the distributions representing the average axon and dendrite of participating neurons must be updated with that information. B4: This can lead to an increased or decreased overlap for connections to and from all other neurons.

While these aspects are not or only incompletely recreated in simple models based solely on evaluating spatially structured connection probabilities, they are found in a model of connectivity predicted from individual reconstructed neuron morphologies [49; 47; 25]. However, previous work on this topic has remained observational, and no explanation has been brought forward to explain this phenomenon. Considering the results closely, we found that all the aspects outlined in Fig. 1A describe types of statistical dependence between connections. That is, the presence of a connection between one pair of neurons increases the probability of a connection between another pair. Based on this, we formulate a hypothesis of how structure emerges when individual morphologies are used and why it does not emerge (or less so) when average neurite shapes are considered.

#### Hypothesis

1. A required condition for the formation of a connection is proximity of the axon to the dendrite.
2. This condition can be approximated by a probability function on the offset between somata of a neuron pair, whose shape is determined by the overall average shape of the dendrites and axons of the classes of neurons considered (Fig. 1B1, B2).
3. However, once a connection at a given distance and direction has been confirmed for a given pre-/post-synaptic neuron, this function must be updated for all its future potential connections. This is because presence of the connection demonstrates that the axon / dendrites are more likely to be oriented towards the point where the connection has been formed (Fig. 1B3, B4). This introduces statistical dependence between connections that cannot be captured by models based on statistically independent evaluations of connection probabilities, even if the shape of the probability function is complex.
4. Such statistical dependencies are directly related to non-random higher order structure, such as the observed overexpression of certain connectivity motifs. Intuitively, if a motif is overexpressed, it means the following: If some of its connections are present in a group of neurons, then the probability that the remaining ones are also present, is increased.

Point (1) is the theoretical basis of using morphological constraints to predict connectivity. Point (2) is a commonly used approximation [28]. Point (3), is the crucial part of our hypothesis. Point (4) describes how the proposed mechanism of point (3) is relevant.

While the hypothesis may be intuitive, we have to demonstrate that its postulated effect can affect the structure of connectivity in measurable ways. To that end, we derived from point (3) predictions about the statistical structure of dependencies between individual connections. Note that the strong type- and domain-preference of inhibitory connectivity [50] has been shown to require additional biases in morphology-based models [45], hence we limit our tests to excitatory connectivity.

### 2.2 Testing the hypothesis in excitatory connectomes

If the mechanism of point (3) affects connectivity, then the following should be true: If a connection exists from a neuron *i* to a neuron, then the probability that another connection exists from *i* to the *nearest neighbor* of *j* is increased over its baseline value. The same holds for incoming instead of outgoing connections of neuron *i*. We tested the prediction in a network of potential connections predicted from rat neuron morphologies [45]. Note these are *potential* connections and hence much more dense than in biology (see Methods). The results confirmed our prediction (Fig. 2A1). For connections beyond 250*µm* the overall probability of a potential connection was below 20%, while it was almost 40% if the nearest neighbor is connected. The proposed mechanism is that the presence of the nearest neighbor connection constrains the distribution of neurite mass to have large values around the location of the neighbor. We found that the effect was stronger for outgoing than for incoming connections, indicating that the constraints were stronger for axons than for dendrites. Additionally, the effect was weaker closer to the soma where the baseline connection probability was already high. We note that half of all potential connections were found at distances beyond 250*µm* where the effect was strong. While these results are based on a single network instance, the connection probability estimates are based on very large numbers of neuron pairs. This lead to highly significant differences between overall connection probability and connection probability for connected nearest neighbors (Fig. 2A1, insets). For a distance-dependent control, as expected, no difference was observed and no statistical significance reached (Fig. 2B1).

**Figure 2:**
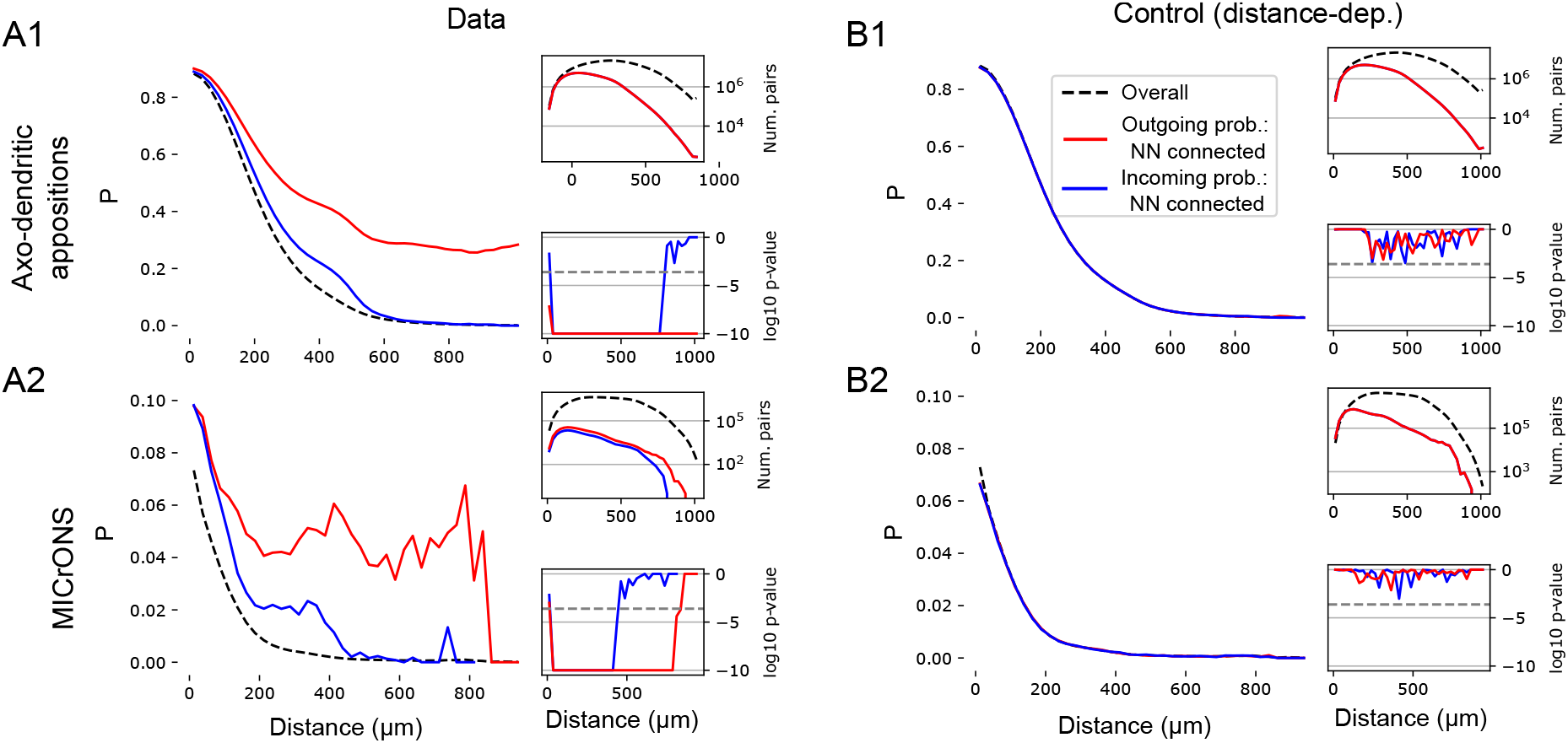
Connections onto a neuron are more likely when its nearest neighbor is also connected A: We test our hypothesis using the connection probability conditioned on the nearest neighbor of a neuron (*NN*) being connected, i.e., *P*(*A* →*B* | *A* → *NN*(*B*)) and *P*(*A* → *B* | *NN*(*A*) → *B*) A: Connection probabilities: overall and for connected nearest neighbors (25*µm* bins). A1: In a model network of potential connections from axo-dendritic appositions. A2: In an electron-microscopic reconstructed network. Insets, top: The number of pairs evaluated for connection probability estimates. Insets, bottom: Logarithm of p-values of a test against the null hypothesis that the connection probability conditioned on a connected nearest neighbor equals the overall connection probability. Grey dashed line indicates significance threshold at p=0.01, Bonferroni-corrected (number of distance bins tested). B, as A, but for for respective distance-dependent controls.

These results confirmed that the proposed effect affects potential synaptic connectivity, however, it remained to be shown that an actual connectome would be affected as well. It has been shown that only between 5% and 10% of potential connections are active in a biological neuronal network and it is possible that in this small subset the effect is degraded and insignificant. Therefore, we repeated the analysis for the excitatory connectome of the MICrONS dataset [10], a dense electron microscopic reconstructions of a volume of mouse cortical tissue. Specifically, we analyzed the adjacency matrix of connectivity between excitatory neurons in the volume. To ensure accuracy of the results, we limited our analyses to 1680 pre-synaptic neurons that were successfully proofread for integrity of their axon. On the post-synaptic side no such filtering was applied and all 45598 excitatory neurons were considered, as inputs to neurons are generally accurate without proofreading of the dendrite [51]. See STAR Methods for exact rules on what pairs were or were not analyzed. Note that the density of connections in this dataset was indeed 20 times lower than for the model used above. We found that our prediction was also confirmed for the MICrONS connectome (Fig. 2A2). While fewer pairs were sampled than for the modeled network, the differences we still highly statistically significant for distances up to 400*µm* (incoming) and 800*µm* (outgoing connectivity; Fig. 2A2, insets). Importantly, we also confirmed a weaker effect for incoming than outgoing connections and relatively weaker closer to the soma, as well as absence of any effect in a distance-dependent control (Fig. 2B2). Taken together, we concluded that point (3) of our hypothesis is confirmed with the proposed effect affecting the structure of both potential and actual connectivity.

Next, we investigated how our results relate to previous work, which shows that much of the connectivity structure is driven by the shape of the degree distributions. In particular, it is known that neuronal degree-distributions are long-tailed and this creates structured connectivity [31]. In line with our arguments, Piazza et al. [52] argue that this is a consequence of axon physicality and they describe a multiplicative axon growth mechanism that gives rise to a log-normal distribution of axon lengths. In this light, we have to consider the following explanation: Once a connection to the nearest neighbor has been confirmed, it is more likely that its axon length comes from the long tail of the distribution, making all future connections more likely. This would result in a global connection probability increase for that neuron to all others, not limited to the nearest neighbor. Conversely, the effect outlined in Fig. 1B is based on the axon being confirmed to be present near an innervated neuron leading to a more localized increase of connection probability in its neighborhood.

To tell the two alternatives apart we repeated our analysis, but instead of distance bins we considered two-dimensional spatial bins. Specifically, 50*µm* bins of the offset along the vertical axis (orthogonal to cortical layer boundaries) and horizontal distance (Fig. 3A). This allowed us to assess the spatial structure of the increase in connection probability better. Additionally, we repeated the analysis for a *configuration model* control of the data. This is a stochastic control that shuffles the locations of connections, but preserves the in- and out-degrees of neurons, thereby capturing the effect of long-tailed degree distributions. We found that the nearest-neighbor-based increase in connection probability strongly depended on the location of the spatial bin considered (Fig. 3B, C). This spatial dependency was stronger for the biological connectome than for the modeled one. The spatial dependence cannot be explained by the long-tailed degree distributions. This was confirmed by the results for the configuration model controls, which had a much weaker, spatially relatively uniform increase in connection probabilities when nearest neighbors are connected (Fig. 3D, E right). Again, for incoming connectivity the increase was substantially weaker (Fig. 3D, E left). We conclude that the nearest-neighbor-based increase in connection probability is a combination of two factors: a global increase resulting from long-tailed degree distributions, and a local, spatially constrained increase. Piazza et al. [52] have pointed out that the degree distributions are a consequence of axon physicality and a multiplicative axon growth mechanism. The local increase results when an axon is confirmed to be physically present in a given neighborhood. Hence we will refer to these factors as the “global and the local part of the axon physicality effect on neuronal connectivity”. However, we also point out that they are two parts of the same effect and that the distinction can be argued to be artificial.

**Figure 3:**
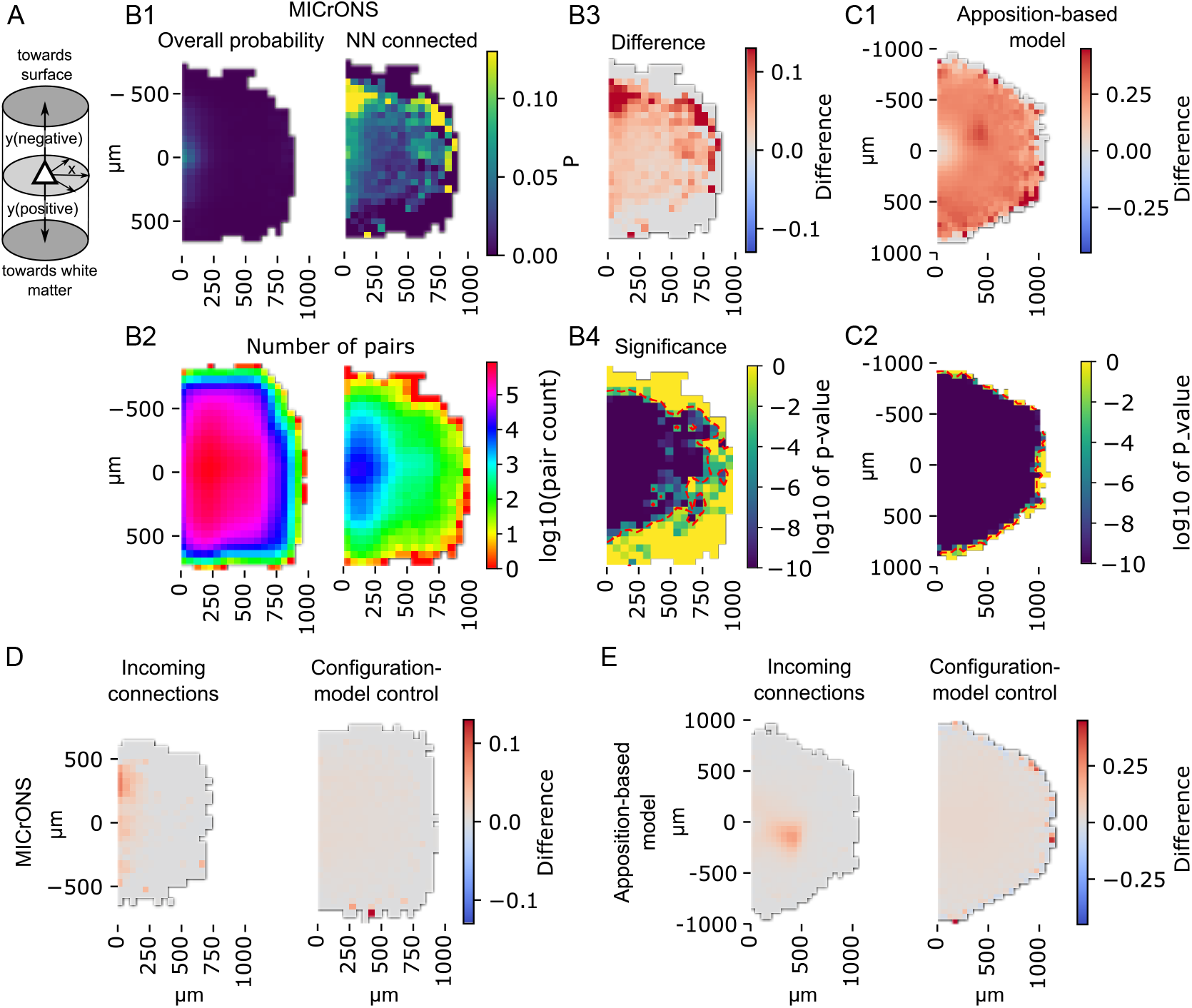
The effect of the nearest-neighbor on connection probability has a complex spatial structure A: Coordinate system for a 2-dimensional version of the analysis in Fig. 2: y-axis indicates the vertical offset of a potential connection (i.e., pair of neurons) with negative values for connections towards the cortical surface, positive values towards the white matter. x-axis indicates the horizontal offset in any direction orthogonal to the y-axis. B1 left: Overall outgoing connection probability in the MICrONS data against horizontal and vertical soma offset (50*µm* bin size). Right: Same, but for nearest neighbors confirmed connected. B2: Number of pairs evaluated for the connection probabilities in each bin indicated in B1. B3: Difference between B1 left and right. B4: Logarithm of p-values of a test against the null hypothesis of equal connection probabilities. Red contour: Area of statistical significance (*p <* 0.01, Bonferoni-corrected against number of spatial bins tested). C1, C2: As B3, B4 but for potential connectivity predicted from appositions. D, left: As B3, but for incoming connections. Right: As B3, but for a configuration-model of the data (degree-preserving control). E: As D, but for potential connectivity from appositions.

So far, our analyses captured statistical dependencies between connections to a neuron and to its nearest neighbor. The effect proposed in our hypothesis could potentially have longer-reaching effects, reaching further into the neighborhood of a connected neuron. To investigate its reach, we conducted the following analysis. We considered for each individual neuron its connection probability into the same spatial bins as before. Then, we calculated the Pearson correlation of connection probabilities of pairs of bins over neurons. A positive correlation does not indicate that connection probabilities are similar, but that a pair of bins is more likely to be innervated together than expected from their individual connection probabilities. A negative value would indicate that their innervation is to some degree mutually exclusive.

We found strong correlations of MICrONS connection probabilities between spatial bins (Fig. 4A; figure shows results for two exemplary bins; see STAR Methods for instructions how to obtain results for any bin). In general, positive correlations up to 0.7 were found for spatial bins within 250*µm* of another bin, although the exact shape of the correlation structure was more complex and not always circular around a bin. Correlations were stronger and extended further for upwards than for downwards connections, in line with Fig. 3B3. Notably, we observed by far more and stronger positive than negative correlations, and statistical significance was reached only for positive ones (Fig. 4E). A negative correlation could be interpreted as follows: If an axon innervates one given spatial bin, it has less remaining “energy” to innervate other bins. This would be in contradiction with Piazza et al. [52], where continued growth of individual axon branches is independent of each other. Hence, our result supports their ideas. For a distance-dependent control, correlations were close to zero, as expected (not shown).

**Figure 4:**
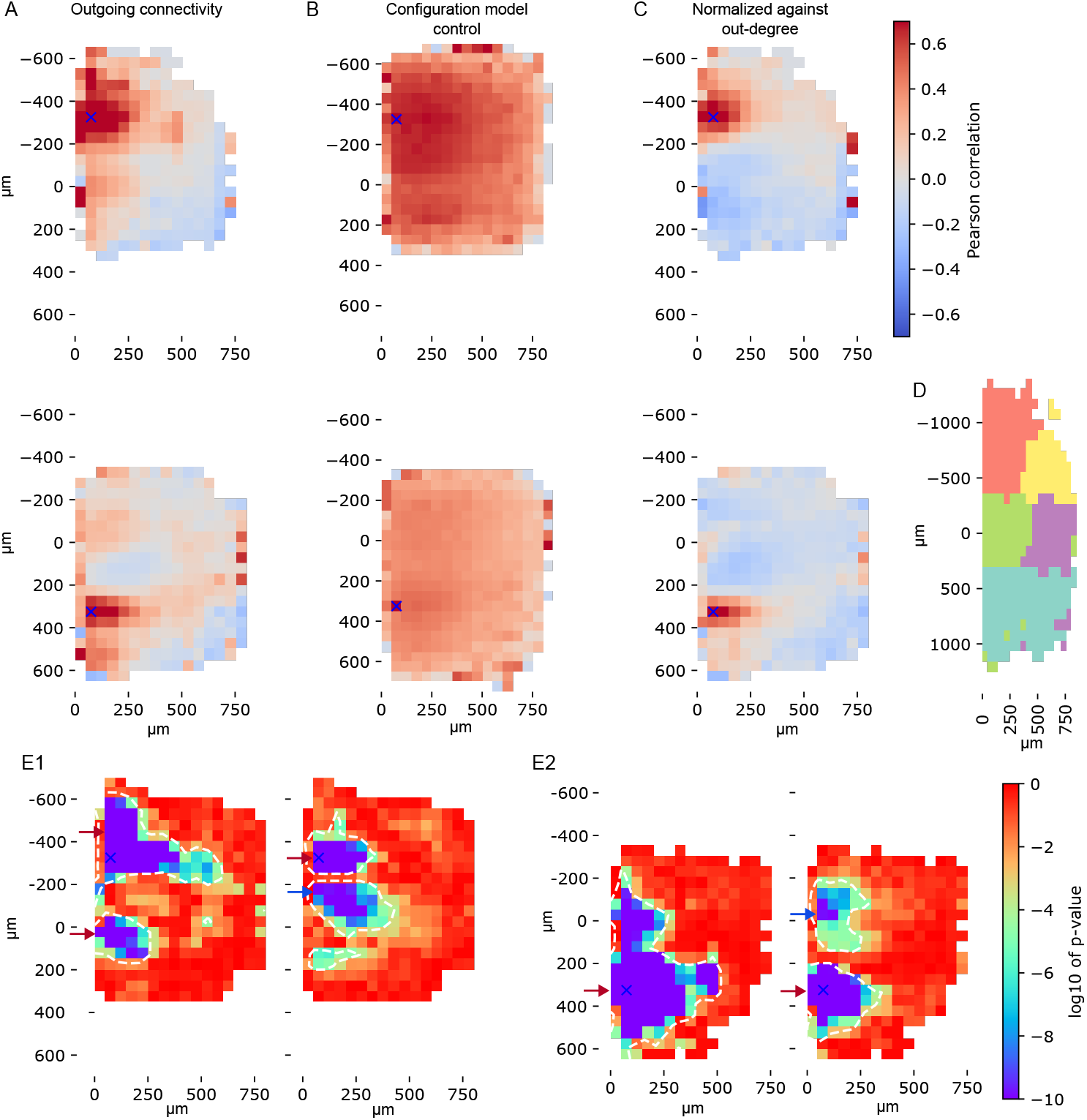
Correlations between innervation strengths into spatial bins. Note that this figure uses the same coordinate system that is illustrated in Fig. 3A. A: Top: Pearson correlation over neurons of outgoing connection probabilities into a spatial bin at a vertical (orthogonal to cortical layers) offset of 325*µm* and a horizontal offset of 75*µm* (blue x-mark) and all other spatial bins. Bottom: As top, but at a vertical offset − 325*µm* instead. Negative vertical offsets indicate connectivity towards more superficial layers, positive values towards deeper layers. B: As A, but for a configuration model control. C: As A, but after normalization for differences in out-degrees. D: Louvain clusters of the matrix of correlations after normalization (i.e., as in C). Correlations *<* 0 were set to 0. E: Logarithm of p-values of correlations depicted in A and C. E1: For the data in the top row before (i.e., as in A, left) and after (i.e., as in C, right) normalization. E2: Same, for the data in the bottom row. White contour: Area of statistical significance (*p <* 0.01, Bonferoni-corrected against number of spatial bins tested). Note that before normalization significance is reached only for regions with positive correlations (red arrows), while after normalization regions of significance are pairs of positive and mostly negative (blue arrows) correlation.

For the configuration model control results were again largely global rather than spatially structured, as expected (Fig. 4B). Remaining spatial structure was the result of inhomogeneous neuron densities in the MICrONS data and dependence of degrees on depth. We confirmed this by analyzing a configuration model where additionally neuron locations were randomized to uniform density (Fig. S1 A, B). Curiously, correlations for the configuration model were also stronger than for the data in many spatial bins. This led to the prediction that the local effect leads to negative correlations that cancel out the positive ones induced by the global effect,i.e., by long-tailed out-degree distributions. To try to capture this, we normalized the connection probabilities estimated for individual neurons: We divided the estimates for individual spatial bins by the mean for a neuron over all spatial bins. This removes the trend of neurons with a larger out-degree having higher connection probability estimates, and thus the effect of long-tailed degree distributions. We confirmed the efficacy of this normalization by confirming that it reduces correlations observed for the configuration model to close to zero (Fig. S1 C). For the MICrONS data, correlations with the global effect thusly removed had regions of statistically strongly significance negative correlation (Fig. 4E). For incoming connections, resulting correlations were consistent with the observations above, but once again significantly weaker (not shown). For potential connectivity predicted from morphologies we found largely the same trends (Fig. S2). While some negative correlations were visible before normalization, they were less abundant and several times weaker than the positive ones. Overall correlations were stronger; however, we note that large differences in sparsity between the connectomes makes quantitative comparisons difficult.

To better understand the structure of the correlations, we ran a clustering algorithm [53] on the matrix of pairwise correlations for outgoing connections after normalization. The clusters obtained were groups of spatial bins that tended to be innervated together (Fig. 4D). We found one cluster for upwards connections with low horizontal distances (red), one for upwards connections with higher horizontal distances (yellow), one for downwards or horizontal connections with distances below 400*µm* (green), and one each for downwards (teal) and for horizontal connections (purple) beyond 400*µm*.

In summary, we further confirmed that neurite physicality affects connectivity by introducing statistical dependencies between connections. We also found that the proposed effect can be described by a combination of a global connection probability increase that is captured by the long-tailed degree distributions commonly found in biological neuronal networks and localized increases and decreases that have a complex spatial structure.

### 2.3 A simplified model of the axon physicality effect leads to non-random micro-structure

Thus far we have demonstrated that the effect related to neuron physicality proposed in point (3) of our hypothesis affects potential and actual connectivity. Point (4) explains why this could lead to the non-random structure observed in biological neuronal networks. However, it is still possible that features such as overexpression of reciprocity, overexpression of specific connectivity motifs [17; 26; 18] and non-random locations of reciprocal connections [25] are the result of other mechanisms. In this section, we will demonstrate that the morphology effect explored above is capable of generating all these features. To this end, we developed an algorithm for the stochastic generation of a connectome graph that is not based on neuron morphology, but still captures its effect of inducing stochastic dependencies between connections.

We call our model for local neuronal wiring a *stochastic geometric spread graph (SGSG)*. These are graphs where edges from a source node to target nodes are iteratively placed using a stochastic process that spreads along the edges of an initial graph on the same nodes. The initial graph is a *random geometric graph*, i.e. a graph where each node is connected to a subset of the nodes in its immediate neighborhood. The spread starts from a given source node *i*, iteratively moving along the edges of the random geometric graph, and all nodes reached from it by the stochastic process will be directly innervated by *i* in the model (Fig. 5A). Repeating this for all possible source nodes yields the full graph of the model. Intuitively, the spread along the edges of the random geometric graph mimics the spread of a growing axon through space. This captures the localized increase in connection probability resulting from point (3), i.e., the localized part of the axon physicality effect, for the following reason: In order for a connection between distant nodes (such as 0 and 11 in Fig. 5A) to exist, the stochastic process has to spread along multiple edges with the possibility of failure each time. However, once it has reached a neighboring node (e.g. node 8), only a single additional spread is required, greatly increasing the probability.

**Figure 5:**
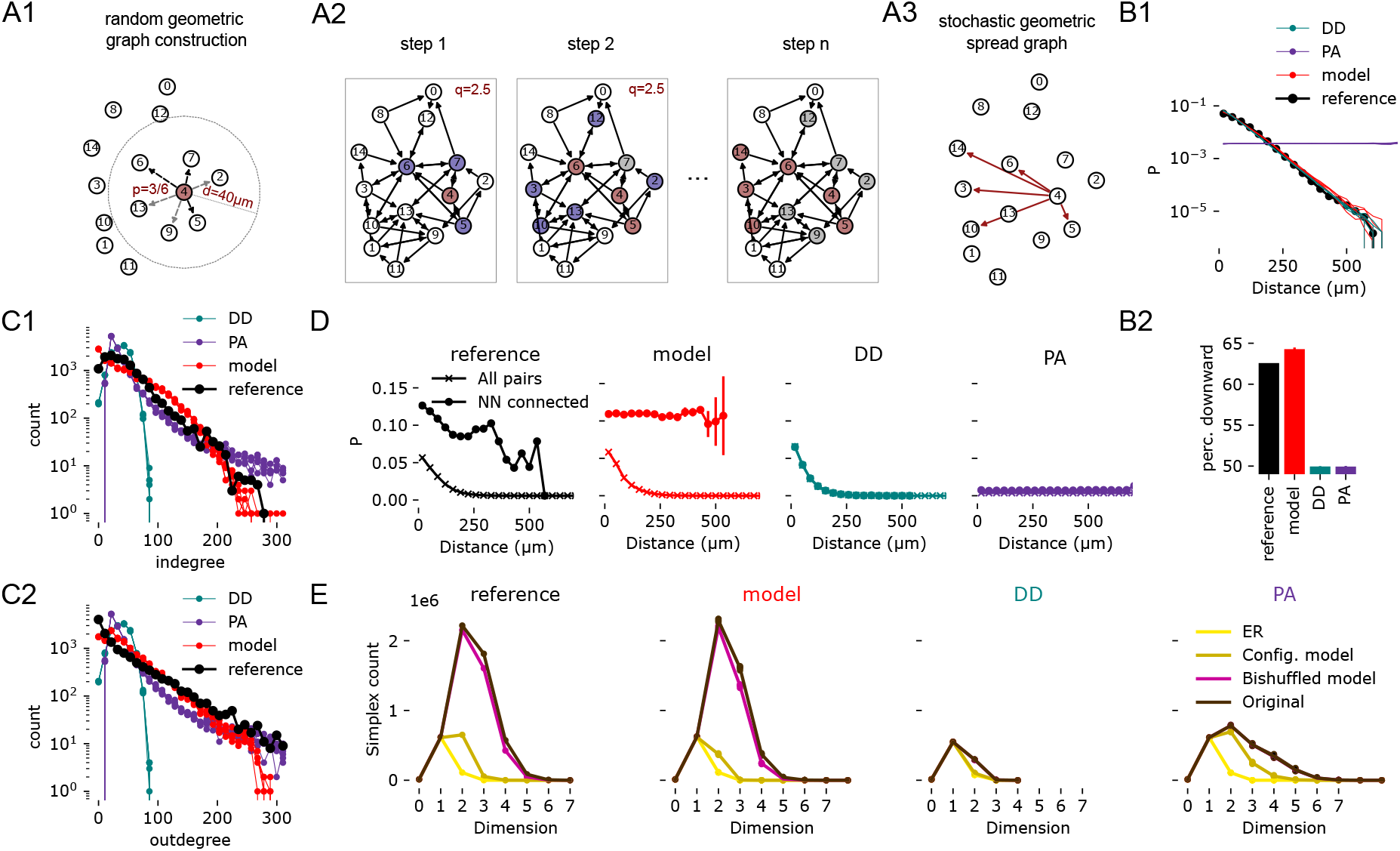
A model of the effect of neuron physicality (SGSG) recreates biological non-random connectivity. A: Construction of a stochastic graph model (SGSG). Example for node 4. A1: A node is associated with a location and other nodes within distance *d* are detected (grey). A directed edge connects them (black) with probability *p* (random geometric graph). A2: For a given node (red) the spread process selects its neighborhood (blue) as candidates. Nodes are selected from them randomly with an expected number of *q* nodes selected. Step 2: The neighborhood of the newly selected nodes (red) comprises the new set of candidates (blue), but nodes than were previous candidates but not selected (grey) are excluded. Step *n*: The process terminated after no new nodes have been selected. A3: In the SGSG, an edge exists from the starting node to all nodes reached by the process. B-E: Results after fitting against a reference network (MICrONS) with meso-scale additions to the algorithm (“model”, see Methods), as well as results for distance-dependent (DD, green) and preferential attachment (PA, purple) models fitted to the reference. For model, DD and PA results of five instances shown. In panels B2, D: mean ± standard deviation, in all other panels: individual instances are shown. B1: Distance-dependence of connectivity for reference, model and controls. B2: Percentage of downwards connections. C: Distribution of in- (top) and out- (bottom) degrees. D: Connection probability increase if nearest neighbor is connected (as in Fig. 2). E: Counts of directed simplex motifs in the respective data and controls of increasing complexity (see Methods). Note that for model, DD and PA individual instances are shown, but overlap completely.

The SGSG model has three parameters, two of them (*d, p*) determine the underlying random geometric graph and one (*q*) parameterizes the spread process. *d* is the maximum distance of connection in the random geometric graph; *p*, the probability that any of the pairs within that distance are connected. *q* parameterizes how likely it is that the spread process crosses an edge of the random geometric graph: In each step, first the number of edges that can be crossed is calculated, then the probability of crossing is set to *q* divided by that number. For a full, mathematical description of parameters and algorithm refer to the STAR Methods. We began by evaluating this family of graphs with respect to several features of connectivity, scanning over a wide range of parameter values (Supplementary section S3, Fig. S9). Briefly, we found that SGSG have: (1) distance-dependent connectivity with different degrees of steepness of decrease for different parameter combinations; (2) long-tailed (lognormal) degree distributions with different degrees of skewness; (3) non-random microstructure in forms that have been characterized in biological neuronal networks, i.e., clustered connectivity, overexpression of directed simplices and reciprocal connections. Importantly, the long-tailed degree distributions meant that also the global part of the axon physicality effect can be captured by the model. We conclude that SGSG, a simple graph model based on the effect outlined in point (3) of our hypothesis, does indeed generate various relevant features of non-random structure. This supports point (4) of our hypothesis. Additionally, we note that the SGSG model is versatile, capable of generating networks with very different characteristics at different locations of its parameter space. In fact, for *d* → ∞, *p* = 1 the model is equal to a model with a single uniform connection probability (Erdos-Renyi graph) with density 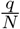, where *N* is the number of nodes.

While the above demonstrated the presence of non-random connectivity features in SGSGs, it remains unclear if they are capable of generating them to the same degree as found in biological neuronal networks. To test this, we fitted an SGSG to the connectivity of excitatory neurons in layers 4 and 5 of the MICrONS connectome (from here on: the “reference”). The first step was to use the soma locations of the reference in the construction of the random geometric graph. In addition to non-random *micro*-structure, biological neuronal networks also have a characteristic *meso*-structure. To improve the meso-scale match, we used a number of additional customizations of the SGSG model. Connection probability in the reference decreased more slowly along the y-axis (roughly orthogonal to layer boundaries), indicating a columnar organization of connectivity [54]. To match this, we simply divided the y-coordinates of neurons by a factor *f*_*y*_ for the construction of the random geometric graph, thereby decreasing distances along that axis. Furthermore, we added an *orientation bias, w*_*A*_, of connectivity. Specifically, we made downward connections (towards deeper layers) more likely in the random geo-metric graph, and those going upwards less likely. Finally, we added *per node biases, w*_*o*_ and *w*_*i*_, that made connections from (*w*_*o*_) and to (*w*_*i*_) nodes more or less likely. These were respectively calculated based on the mean outgoing and incoming connection probabilities of the morphological type a neuron belonged to. Hence, this introduced parameters, but they were deterministically calculated and not subject to a fit. Note that the three customizations affected the random geometric graph (Fig. 5A1) and not the spread mechanism (Fig. 5A2).

We were able to obtain a match to the reference for many relevant aspects of connectivity. Connectivity decreased with distance linearly in log-space (Fig. 5B1), with a matching spatial bias making downwards-facing connections more likely (Fig. 5B2). Degree counts were long-tailed with a steeper slope for out-degrees, unlike in a distance-dependent control (Fig. 5C). We confirmed a match for increased connectivity when the nearest neighbor is connected (Fig. 5D), the core mechanism of our hypothesis. For counts of directed simplices we also obtained a near-perfect match to the reference (Fig. 5E). Coarse-grained pathway strengths were reasonably well matched (Fig. S3). Alternative models of local connectivity did not match the reference in all aspects. A distance-dependent (DD) model matched - unsurprisingly - the distance profile, but not the degree distributions, while for a preferential attachment (PA) model it was the other way around. However, both DD and PA did not exhibit the strong increase in connection probability for connected nearest neighbors. A small, but significantly weaker increase was observed for PA (see also Fig. S4). For simplex counts, again the profiles of both DD and PA were more distant to the reference than SGSG. Additionally, only the SGSG matched a trend found for the reference that simplex counts were higher than in a “bishuffled control” (Fig. 5E, black vs. purple). That control maintains most of the topology of the reference network and only randomizes the locations of bidirectional edges. This indicates that the locations of bidirectional edges are non-random with respect to the simplicial structure, which has been previously shown to be the case in biological connectomes [25]. It must be noted that different forms of PA models exist that may provide better matches of simplex counts, however, fitting the models to simplex counts is known to be difficult [55; 56], making the close match for the SGSG more remarkable. See Supplementary section S4 for an exploration of additional models.

We repeated these analyses for two additional reference connectomes. For layers 2/3 of the MICrONS data we did not re-fit parameters *d* and *p* to test generalization (Fig. S5; For layers 4/5 of the model of circuitry of rodent somatosensory region of Reimann et al. [45] we did a complete re-fit (Fig. S6. Again, we obtained reasonable matches in measured aspects.

### 2.4 Using the model to describe long-range connectivity

We have shown that our proposed mechanism can shape connectivity in way that recreates relevant non-random structure matching biology. However, thus far this is limited to local connectivity, while most of the connections in the brain are non-local, long-range connections [57]. Here, we sketch out an extension of SGSGs for long-range connectivity. Briefly, we use the union of two random geometric graphs instead of a single. While the second one is built on the same nodes as the first, it uses different node locations (**P**^∼^) in its construction that encode the structure of inter-regional connectivity by proximity. Here, we explore a simple example (Fig. 6A): In an elongated point cloud we draw a region border simply as a dividing plane. One coordinate of **P**^∼^ is then the distance from that line. This will place nodes on opposite sides but at the same distance from the plane close to each other, leading to edges between them in the random geometric graph that are not present when the regular locations (**P**) are used. This approach is based on topographical mappings of connections between primary and higher visual areas that follow such a rule in mouse.

**Figure 6:**
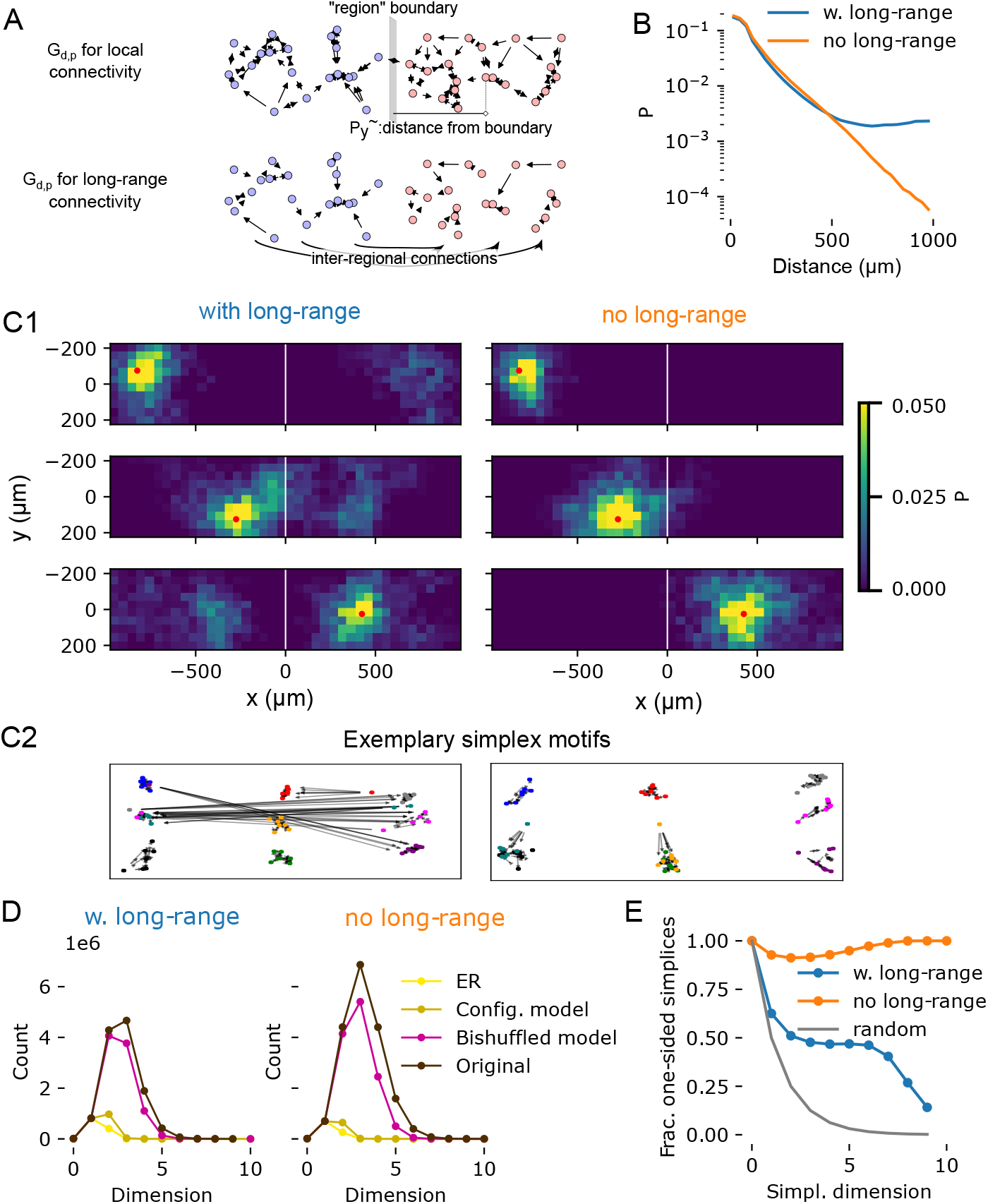
An extension of the SGSG model to long-range connectivity. A: We extent the model by building two random geometric graphs, one for local and one for long-range connectivity, then using the union of their edges to build a stochastic spread graph. The long-range portion uses the same parameters, but “virtual” node locations instead of the real ones. Here, we use virtual locations based on the distance from a region border drawn in the middle of the points. B: Distance-dependent connection probabilities of the resulting SGSGs. Blue: As described in panel A; orange: built without long-range portion, as in Fig. 1. C1: Two-dimensional connection probabilities of neurons in various spatial bins. Red dot indicates the location of a spatial bin, colors indicate the connection probabilities of nodes in that bin to all other bins. White line is the location of the region border. Left column: For three exemplary bins in a connectome model with long-range connectivity. Right column: For the same bins in a model without long-range connectivity. C2: Locations of neurons in exemplary simplex motifs (colored circles) and the connectivity between them (arrows). The model with long-range connectivity has some simplices with neurons on both sides of the region border. D: Simplex counts as in Fig. 2F with and without long-range connectivity in the model. E: Fraction of simplices that do not contain nodes on both sides of the region border. X-axis indicates the dimension of simplices considered. Blue and orange as in B. The grey line indicates results if the side of the border is shuffled for all nodes.

We built the stochastic spread model on such a union of two graphs and contrast it with a stochastic spread graph on only the random geometric graph representing local connectivity. We found that, as expected, the connection probability was largely identical to the graph with only local connectivity below 500*µm*, but beyond that it flattened out, indicating the presence of long-range connections (Fig. 6B). These were also clearly visible, with topographical mapping in the two-dimensional connection probabilities (Fig. 6C1). Consequently, directed simplices, tightly connected and functionally relevant [25] motifs were formed that included the long-range connections (Fig. 6C2). We investigated this aspect further, counting the number of simplices with and without long-range connectivity, and what fraction contains neurons on both sides of the region border. We found the overall number of simplices to be slightly lower with long-range connections in place (Fig. 6D). However, the simplices were much more likely to contain neurons from both sides of the region divide when long-range connectivity was present (Fig. 6E). In fact, large simplices were almost completely limited to only one side without it. As simplices have been shown to be motifs with functional implications [41; 25], this demonstrates that the addition of long-range connectivity in such a model enhances inter-regional interactions.

## 3 Discussion

Non-random micro-structure has been previously demonstrated to emerge in network models built from neuron morphological constraints, and here we have formulated and characterized the mechanism leading to this. To support our claims, we have derived a prediction about the structure of biological neuronal connectivity from our hypothesis, demonstrated that it is not true in simple network models, but that it is true in models derived from appositions between individual neuron morphologies. We also confirmed it in an experimentally measured connectome, demonstrating that the mechanism affects biological connectivity. We have further shown that a stochastic algorithm implementing the proposed mechanism without use of neuron morphologies gives rise to multiple non-random features of connectivity and can be fit to various connectomes. Our proposed mechanism stems from the difference between the average shape of axons over a population of neurons and the individual axons. While the average axon can innervate large parts of the surroundings of a neuron, an individual axon can only reach a small part of it. This part must be spatially continuous due to the physicality of the axon, leading to statistical dependencies of innervation of one neuron and other neurons in its neighborhood. Similar logic can be applied to the shape of the dendrites. However we found that the mechanism plays out much more weakly for incoming connectivity, associated with the dendrites, than for outgoing. The evidence we presented is already strong indication that our hypothesis is true, but for further proof we propose the following experiment: In the MICrONS electron-microscopy volume, detect all potential synapses, i.e., axo-dendritic appositions. Then characterize the network structure resulting from potential connections according to the analyses used in this manuscript and contrast it with the structure of the actual connectome. This will demonstrate to what degree the non-random network structure is constrained already by the shapes of axons and dendrites.

We have further split the proposed mechanism into two conceptual parts: A global effect that is associated with (and explains) long-tailed distributions of out-degrees, and a local one acting on top of it, adding spatially structured dependencies between connections to different neurons.

This places our work in line with Brunel [31] who demonstrated that long-tailed degree distributions give rise to some non-random structure, and with Egas Santander et al. [25] who demonstrated that they do not explain all structure in biological neuronal networks, i.e., that there must be a mechanism acting on top of them. That the axon as a physical object underlies the long-tailed degree distributions has also been pointed out by Piazza et al. [52], who described its growth as a multiplicative process. This is in line with our result that spatially binned correlations of connection probabilities over neurons are positive (unless normalized). Additionally, our stochastic graph algorithm that implements the proposed mechanism also describes a multiplicative process. Also in line with Piazza et al. [52], we found that degree distributions in our model were log-normal when fitted to biological data from cortex (Fig. S9). However, at other locations of the parameter space this may differ. For example, Erdos-Renyi models are a special case of the SGSG model and have binomally distributed degrees. Similar to our ideas, Liu et al. [58] proposed an algorithm for long-range connectivity based on explicit, but abstracted simulations of an axon growing process. However, in their algorithm each axon innervates only a single node, thereby missing the statistical dependencies that we have shown to matter. Others aim to explicitly simulate the process of network formation during development [59]. It should be noted that while our algorithm conceptually grows an axon from the soma this is not intended to capture the actual development, but only the resulting network structure. We have shown that models based on distance-dependence can capture the meso-scale structure of connectivity, and models based on preferential attachment can capture the micro-structure of connectivity, but neither captures both. Proposals exist that combine terms from both approaches into a compromise [34; 35; 36], but in our model these two aspects are inseparably linked into the same mechanism.

Our work should be contrasted with prior research on fractal networks. In particular [37], which introduces a multifractal network generator with tunable parameters that modulate real-world network multifractality. This is complemented by [38], which develops node-based fractal metrics to characterize network complexity and heterogeneity, and by [11], which enables inference of these structures from partial and noisy data. Remarkably, Yin et al. [5] reveals that neuronal culture networks in rodents exhibit unique multifractal and assortative connectivity patterns not captured by classical models. Multifractal network models effectively reproduce generalized degree distributions and clustering coefficients, reflecting higher-order interactions of dimension two. However, how well they capture higher-dimensional structures across all dimensions still remains open.

Moreover, this models predominantly infer network structure through top-down statistical observations, enabling broad applicability across diverse systems. While this versatility extends to data coming from yeast genome networks and functional networks of Alzheimer’s disease patients, it lacks the biophysical spatial constraints specific to microscale neural networks. Our work seeks to address this gap.

In summary, our work builds upon and extends previous work, leading to a unified description of three ways in which neuron morphology structures connectivity: Distance-dependence, derived from the average shape of morphology, degree-distributions, derived from individual neurite lengths, and a statistical dependence, derived from individual neurite shapes.

Our work is relevant because the non-random micro-structure of connectivity is relevant. It has been described in multiple publications, but it origins remained elusive. We have demonstrated that already constraints given by neuronal morphology lay the groundwork for it. It is still likely that additional mechanisms, such as structural synaptic plasticity further reinforce it [60]. This is evidenced by non-random micro-structure in network models built from neuron morphologies being weaker than in biological neuronal networks [61]. Furthermore, a simulation study has predicted that plasticity acts in way that reinforces specifically the non-random simplicial structure of the network [41]. Nevertheless, the link between neuron shape and network micro-structure we established here may be important for our understanding of pathologies associated with degraded neuron morphologies [62; 63; 64; 65] or differences in network structure between species. We note that our results caution against using average neurite shapes for modeling connectivity[28], as that approach will miss the micro-structure. Additionally, our stochastic graph algorithm may help us to understand the functional impact of non-random network structure further, as these structures likely contribute to the higher-order patterns observed in functional networks [66; 67; 68]. This topic has already been studied in biophysically-detailed network models, with some of the predictions confirmed in experimental data [41]. For example, non-random structure has been predicted to be associated with overcoming the lack of reliability of stochastic synapses [25]. Our algorithm allows this to be also addressed in more simplified models of point neurons, as it can wire a network of 20,000 neurons in under 5 seconds on a laptop computer. As such, our algorithm complements studies exploring function-structure relationships using point neuron models with fractal network architectures [69], research in machine learning that examines networks with complex and diverse topologies [70; 71], as well as work that studies how the higher order structure of the network are a driving force for the system achieving self-organized criticality [72; 73]. The algorithm can also play a role for modeling the activity in cultured neuronal networks. In cultures, experimenters can influence the macro-scale structure of connectivity by introducing physical patterns to the environment that guide axon growth, where axons cross the borders of the patterns with lowered probability that depends on the angle of the crossing [74]. Modeling of this setup [75] can make use of SGSG networks with modifications to the underlying random geometric graph (increased distances across borders and orientation bias). Finally, our results encourage the development of sampling strategies for researchers that measure local circuit connectivity experimentally. The strategies should enable the characterization of the statistical dependencies we described and fitting of SGSG models to the data.

Our proposed mechanism is based on the difference between individual and average axons. But the connectomes we have considered contain neurons that are sometimes grouped into different classes. Thus we must ask: Is it the difference between individual and average, or the difference between neuron classes that matters? Our stochastic algorithm generated non-random structure already without different classes of neurons; when we added classes later on this only served to improve the meso-scale structure. Additionally, our hypothesized mechanism would still be relevant in a population of neurons with identical, but randomly rotated axons. This is because axons typically have no rotational symmetry [45].

### 3.1 Limitations of the Study

At the core of many of our arguments is the connectivity of the MICrONS data, which has inaccuracies. For synapse detection, 96% precision and 89% recall is reported with partner assignment accuracy of 98%[10]. Another potential error source are segmentation mistakes, where a neurite subtree and its synapses are incorrectly assigned to a different nearby neuron. Hence we limited our analyses in Figs. 2, 3, 4 to connections where the axon of the pre-synaptic neuron has been successfully proofread. On the post-synaptic side we did no filtering, as segmentation mistakes are less likely for dendrites due to their higher diameter. Additionally, we argue that our results are robust against errors due to the large number of pairs sampled and the extremely low p-values encountered. Finally, we argue that such errors are more likely to destroy than create the structure we found, as a previous analysis of MI-CrONS showed that non-random connectivity is stronger in proofread than non-proofread neurons [61].

While we have shown that the SGSG model can be fit to match connectivity of local cortical circuitry, in its basic form the SGSG model is certainly too simple to capture all aspects of brain networks and their development. We have outlined an extension to long-range connectivity to demonstrate the capability of the model to be improved by extending the notion of the underlying random geometric graph. Here, we want to touch on other missing aspects and outline how they are readily added to the model.

First, while the model is meant to capture the impact of morphology on connectivity, it was fitted without taking actual neuron shapes into account and we have not provided a way to predict morphology from the parameters. While a more systematic study of this is required, we can already speculate: Dendrites extending further away from the soma could lead to a higher ‘d’ parameter. The presence and density of dendritic spines could increase the ‘p’ parameter, effectively increasing the probability the connections are received from locations that dendrites spread to. Additionally, we would argue that the role of dendrite shape could be best captured as follows: A neuron is represented by several nodes in the random geometric graph, one for the soma and a number of nodes along its dendrite. Nodes associated with the dendrite each represent comparable amounts of dendritic surface and have no outgoing edges in the random geometric graph. The incoming edges of a neuron in the final SGSG is the union of edges into its soma- and dendrite-associated edges. This would naturally lead to a higher in-degree for larger neurons and create some degree of asymmetry (such as a laminar structure for cortex) where dendrites are asymmetrical. This can be implemented with a single additional parameter: the amount of dendrite surface per additional node. For perisomatically-targeting Basket Cells [50] the nodes representing distal dendritic domains can be ignored.

Second, the nature of the stochastic spread model implies that every part of an axon forms synaptic connections, as the axon can only spread further from nodes it has formed a connection on. While this is roughly in line with the idea of minimization of wiring cost [76], it is also known that there are stretches of axon with few or no synapses. A potential solution is already outlined by the process in Fig. 6: The construction of a random geometric graph in a custom space where neurons that are distant in the actual brain coordinates are placed next to each other leads to neurons innervating distant patches with little to no innervation on the path to that patch (Fig. 6C). While the given example is very simplistic, more complex spaces could implement more complex mid- to long-range connectivity patterns. For example, connectivity in visual regions has been shown to be structured by similarity of orientation selectivity, which in some animals is structured in repeating patches [77]. We propose to model this by appending preferred orientation as a dimension in addition to the three spatial ones that define the random geometric graph.

Third, while our model generates non-random structure on the per-node level, there is also important structure on the population level: Certain classes of inhibitory neurons have been shown to target other inhibitory neurons [50] in a way that is not explained by neuron morphology without introducing a selection bias [45]. Conversely, in the MICrONS data we based many of our tests on, excitatory neurons have been shown to prefer connections onto inhibitory neurons more than predicted by morphology [10]. We note that for the purpose of this work we avoided these aspects by focusing on excitatory to excitatory connectivity only. We also predict that this structure can be implemented similar to the method of the previous point: By considering dimensions in addition to the spatial ones in the setup of the random geometric graph. In this case, they would define a distance based on molecular neuron classes. Asymmetry in connectivity between classes could be introduced as a directionality bias along the additional dimension, similar to the one we used along the y-axis in (Fig. 5).

In summary, we predict that many features of shortmid- and long-range connectivity can be implemented in the form of more complex coordinate systems than the three spatial brain coordinates for the construction of the random geometric graph, and we propose that further research be conducted to decode that connectomics space. Data to that end is available, for example in the form of transcriptomic atlases [78] and large voxelized connectivity datasets ([79]) which can be subjected to dimensionality reduction [80] Open questions would be, which coordinates to use and their relative scaling. Additionally, they can define additional dimensions for the construction of the random geometric graph or a separate random geometric graph is to be constructed with them that is then merged with the first, as we did above.

## Resource availability

### Lead contact

Requests for further information, source data, and instructions on how to recreate the analyses in this work should be directed to and will be fulfilled by the lead contact, Michael W. Reimann.

### Materials availability

No materials were used in this computational study.

### Data and code availability

#### Code

Analysis of the simplicial structure of connectomes and generation of control models was performed using the publicly available python package “Connectome-analysis”, which is available on the pypi package tracker (https://pypi.org/project/connectome-analysis) and on github (https://github.com/openbraininstitute/connectome-analysis). The implementation of the SGSG model has been added to the same package.

Loading of and analysis of connectome representations was performed using the “Connectome-Utilities” python package available on pypi (https://pypi.org/project/Connectome-Utilities) and github (https://github.com/openbraininstitute/ConnectomeUtilities).

Jupyter notebooks to generate the manuscript figures and instructions how to do so can be found under: http://github.com/MWolfR/local_connectivity_model. This allows generation of Fig. 4 for other exemplary bins.

#### Raw data

The raw data of the MICrONS project can be accessed from: https://www.microns-explorer.org/cortical-mm3.

#### Connectomes

In this work, analyses were run on hdf5-based representations of connectivity data that are optimized for connectomics applications. Specifically, we analyzed four such connectomes: MICrONS in two different versions; and the connectivity of the model of rat somatosensory regions of Reimann et al. [45]. For the later, one version containing all potential synapses, and one version where their number has been algorithmically reduced to a biologically realistic count. Some of the data releases also contain controls of the data (DD=distance-dependent; CM=configuration model), in other cases the controls are calculated ad hoc.

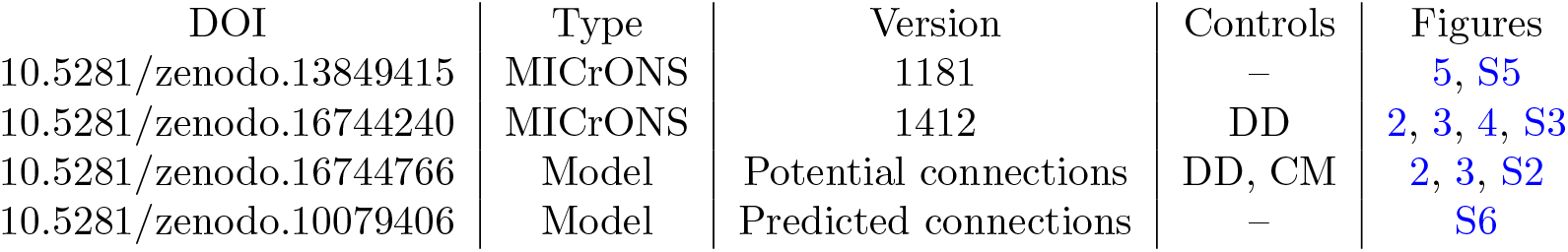

| DOI | Type | Version | Controls | Figures |
| --- | --- | --- | --- | --- |
| 10.5281/zenodo.13849415 | MICrONS | 1181 | – | 5, S5 |
| 10.5281/zenodo.16744240 | MICrONS | 1412 | DD | 2, 3, 4, S3 |
| 10.5281/zenodo.16744766 | Model | Potential connections | DD, CM | 2, 3, S2 |
| 10.5281/zenodo.10079406 | Model | Predicted connections | – | S6 |

## STAR Methods

### Method details

#### Stochastic geometric spread graph model (SGSG)

In this paper, we consider finite directed graphs without loops or double edges. Such a graph *G* is represented by a tuple (*V, E*), where *V* is the set of nodes and *E* ⊆ *V* × *V* is a set of directed edges, with no repeated entries (i.e., (*v, v*) ∈*/ E* for any *v* ∈ *V*).

We define a generative model of random graphs that depend on a single parameter *q*, called the stochastic spread model on a base graph *G* and we denote an instance of such a process by 𝒮_*q*_(*G*). Intuitively, a graph of this type is built from *G* by iteratively spreading from each node of *G* to a subset of its outgoing neighbors, which are selected at random. The size of the subset is also random with an expected value of *q* (Fig. 1A). The process then spreads further into the neighborhood of the newly selected nodes to a new set, and so on. Once a node has been a candidate to spread to once, it will be ignored in all future steps. As the size of the selected subset is random at each step, the empty set is a possible outcome, at that point the process stops. Additionally, the set of excluded nodes grows at each step, also forcing an eventual stop. In the output graph, an edge is placed from the starting node to all nodes reached by the spread. This process is then repeated for all nodes of *G*.

To formally describe this process, let 𝒩_*G*_(*v*) denote the outgoing neighbors of *v* ∈ **V**, i.e., all *w* ∈ **V** such that there is a directed edge in (*v, w*) ∈ **E**. For a subset of nodes *W* ⊆ *V*, we denote by 𝒩_*G*_(*W*) the all the outgoing neighbors of the nodes in *W* i.e., 𝒩_*G*_(*W*) = ⋃_*w*∈*W*_ 𝒩_*G*_(*w*). Furthermore, we denote by 𝒩_*G,r*_(*v*) ⊆ 𝒩_*G*_(*v*) a random subset where each node is selected independently at random with probability *r*. If the base graph is understood from the context, we simply write 𝒩 (*v*), 𝒩 (*W*) and 𝒩_*r*_(*v*) for simplicity.

Now, for each node *v* of *G*, we will recursively define three subsets of nodes, which in step *i* are: the candidate nodes *C*_*i*_(*v*), the selected nodes *S*_*i*_(*v*) and the tested nodes *T*_*i*_(*v*). For *i* = 1 these are given by:

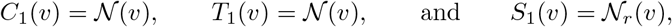

where 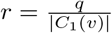. That is, *S*_1_(*v*) is a random subset of the outgoing neighbors of *v* of expected size *q*.

For *i >* 1 we define]

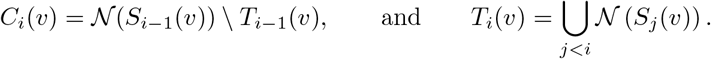

That is, the candidate nodes are the outgoing neighbors of the nodes selected in the previous step which are not nodes that have already been tested in any other step. Then, *S*_*i*_(*v*) ⊆ *C*_*i*_(*v*) is the subset where each node *u* ∈ *C*_*i*_(*v*) is selected independently at random with a probability proportional to the number of neighborhoods it appears in and such that the expected number of nodes selected is *q*. Thus, each node is selected with probability *r* ∗ *m*_*u*_, where *m*_*u*_ is its multiplicity of the appearance of *u* across neighborhoods i.e.,

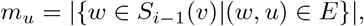

and

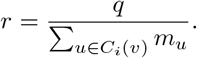

Finally, *S*(*v*) = ⋃_*i*_ *S*_*i*_(*v*) and 𝒮_*q*_(*G*) is a graph with nodes *V* which contains a directed edge from *v* to each node in *S*(*v*) for each *v* ∈ *V*.

At each step, the probability that the process terminates is the probability of obtaining 0 in a binomial distribution with *n*_binom_ equal to the number of candidates for spread and *p*_binom_ = *q/n*_binom_. We can initially approximate *n*_binomm_ ≈ *q* ·|*D*_*G*,out_|, where |*D*_*G*,out_| is the mean out-degree of *G*. It shrinks as the number of nodes excluded from spread grows. Still, we characterize the process as a multiplicative process as described by Piazza et al. [52].

Note that a stochastic spread graph on the empty graph is again the empty graph, while on the fully connected graph on *n* nodes is an Erdos-Renyi graph with overall connection probability 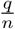.

Additionally, we also consider a type of *random geometric graphs* [81] on a set of points **P** in *n*dimensional space, which we denote by *G*_*d,p*_(**P**). Such a graph has as nodes the set of points in **P** and directed edges added independently at random from *v* to *w* with probability *p* if the euclidean distance between *v* and *w* is smaller than *d*. We call a stochastic spread graph on such a random geometric graph, 𝒮_*q*_(*G*_*d,p*_(**P**)), a <u>stochastic geometric spread graph</u>, which we denote as SGSG.

#### Customizations of SGSGs

We define a number of additional steps to the process that enable customization of the meso-scale or global spatial structure of the resulting SGSG graph. In the context of a connectome model for neuronal circuits, these biases can be used to match neuron type-based or spatial observations in biological connectomes. Briefly, this is done by adding bias terms to the random process selecting which neurons are connected in the random geometric graph. The bias can be based on the types of neurons (per node bias), or the spatial orientation of the potential edge (orientation-based bias).

In order to avoid neurons with zero out-degree, which seems unbiological, we ensure that the first steps of the spread process reach exactly the expected number of neurons (i.e., *q* neurons).

In order to add long-range connections, we consider instead of a singe random geometric graph the union of the edges of two random geometric graphs on the same nodes, but using different node locations.

Details on the customizations above can be found in the Supplementary Methods S2.2.

#### Reference for graph structure - Apposition-based

The reference for connectivity derived from axo-dendritic appositions we based on the network constructed in Reimann et al. [45]. In that work, after detection of axo-dendritic appositions in a population of model neurons, most of the appositions are discarded according to biologically-inspired rules. We instead considered the network generated from all appositions, which we consider the network of all *potential synaptic connections*. This captures the impact of morphology on connectivity without additional synaptic pruning.

#### Reference for graph structure - MICrONS

As a reference for biological neuronal connectivity, we use the wiring diagram for excitatory neurons of the MICrONS connectome [10]. For the analyses related to Figures 1, 2, 3 we used release version 1412 of the data; for Figure 4 version 1181. Only connections between neurons that both had somata inside the volume were considered. For classes of neurons in the data we used the table “aibs metamodel mtypes v661 v2” also provided by the MICrONS initiative [51]. For the excitatory subgraph, we considered neurons where the value of the column “classification system” was “excitatory neuron”.

#### Selecting a subset of connections to analyze in the MICrONS data

While the MICrONS data provides information on the presence or absence of a synaptic connection between all neuron contained in the reconstructed volume, certain inaccuracies are to be expected. A connection is linked to a given neuron by tracing the neurite on the pre- or post-synaptic side to its soma. During the tracing, parts of the neurite of one neuron can incorrectly be merged onto another neuron. This is addressed by manual proofreading of the data, with proofreading annotations provided in the original source.

To compute the connection probability of pyramidal cells in layer 5 in release version 1412 of the data, we analyzed connectivity between 189 neurons classified as one of: “L5NP”, “L5ET”, “L5a”, “L5b”, that passed manual proofreading of both axon and dendrites. The proofreading status is also provided by the MICrONS initiative. Of the neurons, 13,458 pairs were within 100*µm* and 872 of them were connected.

For analyses relating to connection probabilities conditioned on the nearest neighbor being connected (Figs. 2, 3, 4) we only limited our analyses to to neurons with successfully proofread axon. This is because tracing errors are more likely for the thinner axons than for dendrites. Specifically, an ordered neuron pair, (*A, B*) was considered for the calculation of a connection probability *P*(*A* → *B*) only if the axon of *A* was successfully proofread.

The pair was considered for the calculation of a connection probability conditioned on the postsynaptic nearest neighbor being connected if:

1. The axon of *A* was successfully proofread
2. *A* innervates *NN*(*B*)
3. *A* ≠ *NN*(*B*)

The pair was considered for the calculation of a connection probability conditioned on the pre-synaptic nearest neighbor being connected if:

1. The axon of *A* was successfully proofread
2. The axon of *NN*(*A*) was successfully proofread
3. *NN*(*A*) innervates *B*
4. *NN*(*A*) ≠*B*

For the reference for the SGSG model in Fig. 5 and Fig. S5 we considered the entire excitatory subgraph of the indicated portions of the MICrONS data. This is because we only aim to demonstrate that the model can be successfully fit to data with the non-random trends that are typical for biological neuronal networks.

#### Two-dimensional connection probabilities and their statistical dependencies

We calculated for each pair of neurons their offsets in the horizontal plane and along the vertical axis. Offsets were then binned in both directions with a bin size of 50 × 50 *µm*. We refer to a generic bin as 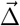.

For the analysis of Fig. 2, we computed the overall outgoing and incoming connection probabilities for any neuron to any other at a given bin. A bin can be regarded as a discretized 2D-vector that represents the offset from the position of one neuron to another. In this way, the analysis provides a two-dimensional analogue of distance-dependent connection probability. We further refined this analysis by considering the conditional probability given that the nearest neighbor is connected. See Supplementary sections S2.1.1 and S2.1.2 for more details.

For Fig. 4, we developed a finer study of this structure by analyzing this probability at the level of individual neurons. Details of this analysis are provided in Supplementary S2.1.3, but we outline the main ideas here.

We fixed a source neuron *v* and considered all neurons either innervated by or innervating it at an offset 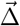, and computed the corresponding outgoing or incoming connection probability. These probabilities can be represented as a vector of length equal to the number of offset bins. We calculated them for all possible source neurons and assembled the result into a matrix with one row per source neuron and one column per offset bin. We calculated the Pearson correlation between pairs of columns of the matrix, resulting in a weighted matrix with rows and columns corresponding to offset bins.

To better understand the structure of this measure, we clustered the resulting matrix. This revealed groups of spatial bins that were more frequently co-innervated by individual neurons (or more frequently co-innervating individual neurons) than would be expected from their global connection probabilities. For all analyses, bins or bin pairs with fewer than 100 data points were excluded.

#### Directed simplex motifs and simplex counts

We counted simplex motifs in the constructed graphs and their controls. A directed *n*-simplex, is a motif on *n* + 1 neurons which are all to all connected in feedforward fashion. That is, there is an ordering of the nodes 0, 1, … *n*, such that there is an edge (*i, j*) whenever *i < j* (see Fig. S7A.). Additional edges are allowed to exists.

#### Graph control models

See also Fig. S7B.

##### Distance-dependent model

First, connection probabilities are calculated in distance bins of size 50*µm*. Then an exponential decay function is fit to the results. For the control, connections are placed randomly with independent probabilities taken from the fitted distance-dependent function.

##### Configuration model

A configuration model is a control that preserves (approximately) the in- and out-degrees of individual neurons. For a reference network, we considered its *edge list*, i.e., a tuple of vectors (*O, I*), with one entry each for each edge. For each index *i* there is an edge from neuron *O*_*i*_ to *I*_*i*_. We shuffled the order of entries of *O*, yielding *O*^*ctrl*^. Next, we removed any entries where 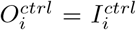 and duplicate entries, i.e., where 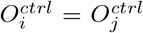 and *I*_*i*_ = *I*_*j*_, we remove entries at index *j*. The configuration model is then the network with edge list (*O*^*ctrl*^, *I*). As the edge list entries are not changed, but merely shuffled, the out- and in-degree of each node are approximately preserved. They will be slightly lowered due to the removal of circular and duplicate entries. Due to the typical sparsity of the networks considered, this loss is minimal.

Notably, the configuration model control also preserves the spatial locations associated with each node/neuron of the reference network. For the analysis in Fig. S1 we additionally assigned new x, y, z coordinates to all nodes. The new coordinates were selected randomly with uniform density within the axis-aligned bounding box of the reference network.

##### Bishuffled model

Most edges are identical, only bidirectional connections are shuffled as follows. For each bidirectional edge a single direction is randomly chosen. Then, in the resulting purely uni-directional graph, a randomly selected subset of edges is made bidirectional to match the number of bidirectional edges in the reference network.

##### Preferential attachment

We used a modified version of the preferential attachment model of Barabási and Albert [29]. In that model, nodes are added one after another and edges are added from each new node to a subset of the previous ones with a preference for high degree nodes. For Fig. 5, we added directionality by randomly selecting a direction for each edge: A bidirectional edge was placed with probability equal to the fraction of bidirectional edges in the reference (MICrONS), otherwise either direction was chosen with equal probability. Other parameters were selected to match node count and mean degree of the reference. In Supplementary Section S4 we also explore the directed PA model of Bollobás et al. [82] and the “small world” network of Watts and Strogatz [32].

### Quantification and statistical analysis

#### Connectivity for connected nearest neighbors

The sampling method explained above resulted for a given connectivity matrix and spatial bin in a list of pairs of neurons such that their soma-to-soma offset falls into the bin, and the nearest neighbor of one of the pair is connected to the other. Let the number of such pairs be *n*. We then test for all pairs whether a direct connection exists between them. Let the number of connected pairs found be *c*. Finally, let the overall (prior) connection probability in that spatial bin be *q*. Our null hypothesis is that the connection probability for connected nearest neighbors is not increased over *p*. In that case, the probability to *c* connected pairs of more is:

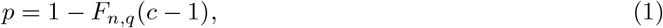

where *F*_*n,q*_ indicates the cumulative binomial distribution with parameters *n* and *q*. Hence, this *p* is the p-value of a one-sided statistical test.

The values of *n* and of 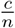 are reported within the figures or in Fig. S8.

#### Significance thresholds

We aimed to use a significance threshold of *p* ≤ 0.01. We performed Bonferroni correction for multiple hypothesis testing by dividing the threshold by the number of spatial bins tested.

#### Correlations between connection probabilities

To assess significance of correlations between connection probabilities into spatial bins (Fig. 4 we used the p-value reported by the “pearsonr” function of the “scipy.stats” python package, version 1.16.1.

The numbers of samples used for this test is reported in Fig. S8.

### Key Resources Table

| REAGENT or RESOURCE | SOURCE | IDENTIFIER |
| --- | --- | --- |
| Deposited data |  |  |
| Adjacency matrix for the MICrONS data, v1181 | [10; 61] | 10.5281/zenodo.13849415 |
| Adjacency matrix for the MICrONS data, v1412 | [10; 61] | 10.5281/zenodo.16744240 |
| Adjacency matrix for potential connections: SSCX model | [45] | 10.5281/zenodo.16744766 |
| Adjacency matrix for predicted connections: SSCX model | [45] | 10.5281/zenodo.10079406 |
| Software and algorithms |  |  |
| Connectome-analysis | [25] | openbraininstitute/connectome-analysis |
| Connectome-Utilities | [45] | 10.5281/zenodo.10059227 |
| SGSG model and analyses | This paper | MWolfR/local_connectivity_model |

## Author contributions

- Conceptualization: MWR
- Methodology: MWR, DES, LK, NBZ
- Software: MWR, DES
- Validation: MWR, DES, LK, NBZ
- Formal analysis: MWR, DES
- Investigation: MWR
- Data curation: MWR
- Writing - Original Draft: MWR
- Writing - Review & Editing: MWR, DES, LK, NBZ
- Visualization: MWR, DES
- Supervision: MWR

## S1 Supplementary Figures

**Figure S1.**
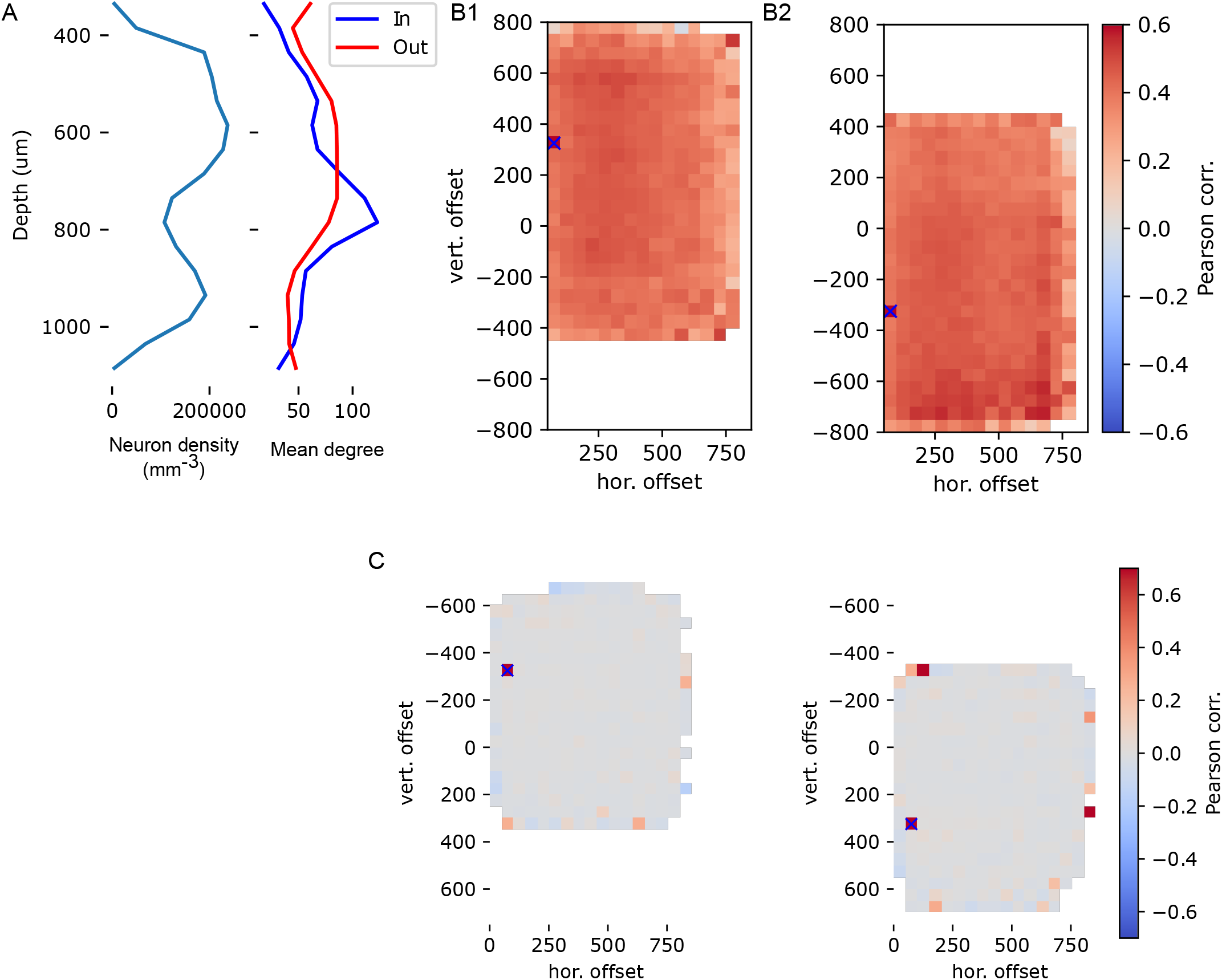
A, left: Neuron density against depth in the part of the MICrONS data we considered in this work. A, right: Mean in- and out-degree of neurons in depth bins in the MICrONS network we considered in this work. B: As Fig. 4B, but for a configuration model control with neuron locations randomized to uniform density. C: As Fig. 4 B, but after normalization against out-degree. That is, as Fig. 4 C, but for the configuration model control. Note that the normalization almost completely erases the correlations.

**Figure S2.**
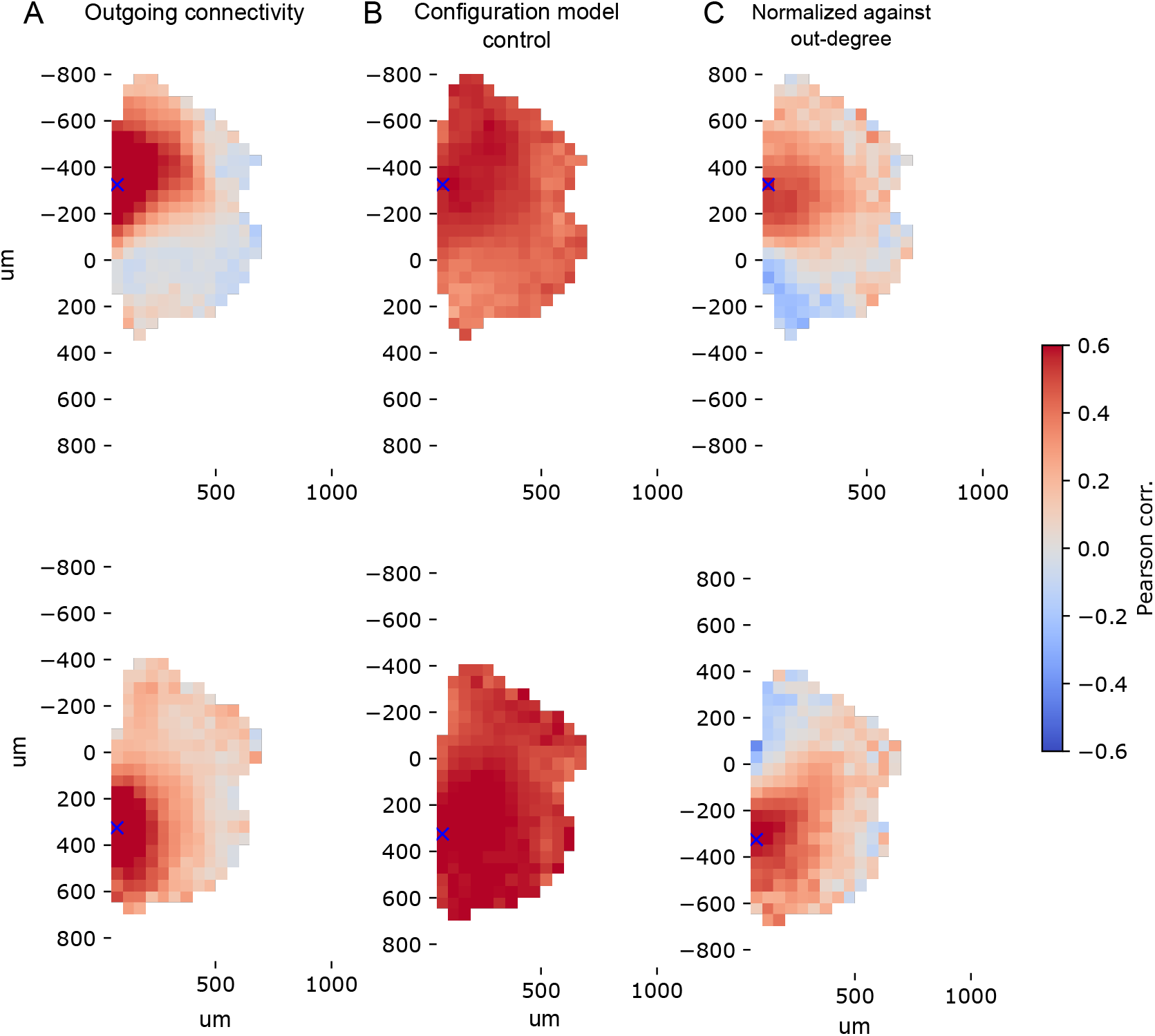
Pearson correlations of connection probabilities into spatial bins as in Fig. 4, but for the apposition-based potential connectome instead of the electron-microscopically measured one.

**Figure S3.**
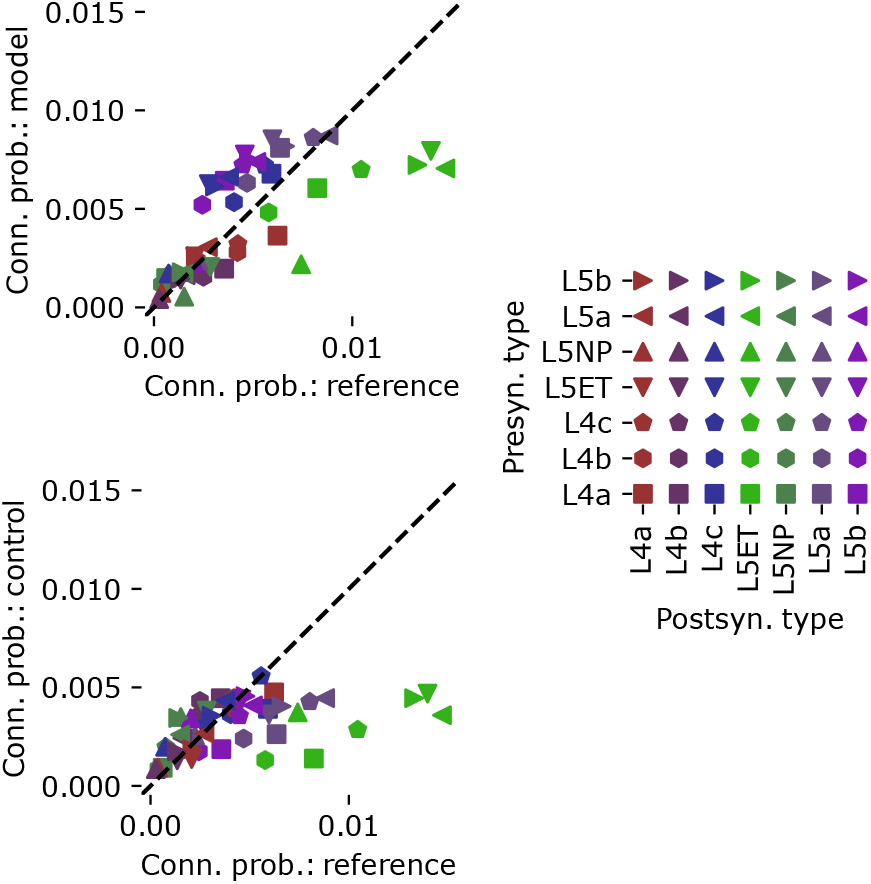
Pathway-specific connection probabilities, reference (MICrONS) against SGSG model (top, pearsonr=0.7) and against distance-dependent control (bottom, pearsonr=0.45). Mean over five instances. Marker shape indicates pre-synaptic, color post-synaptic neuron types.

**Figure S4.**
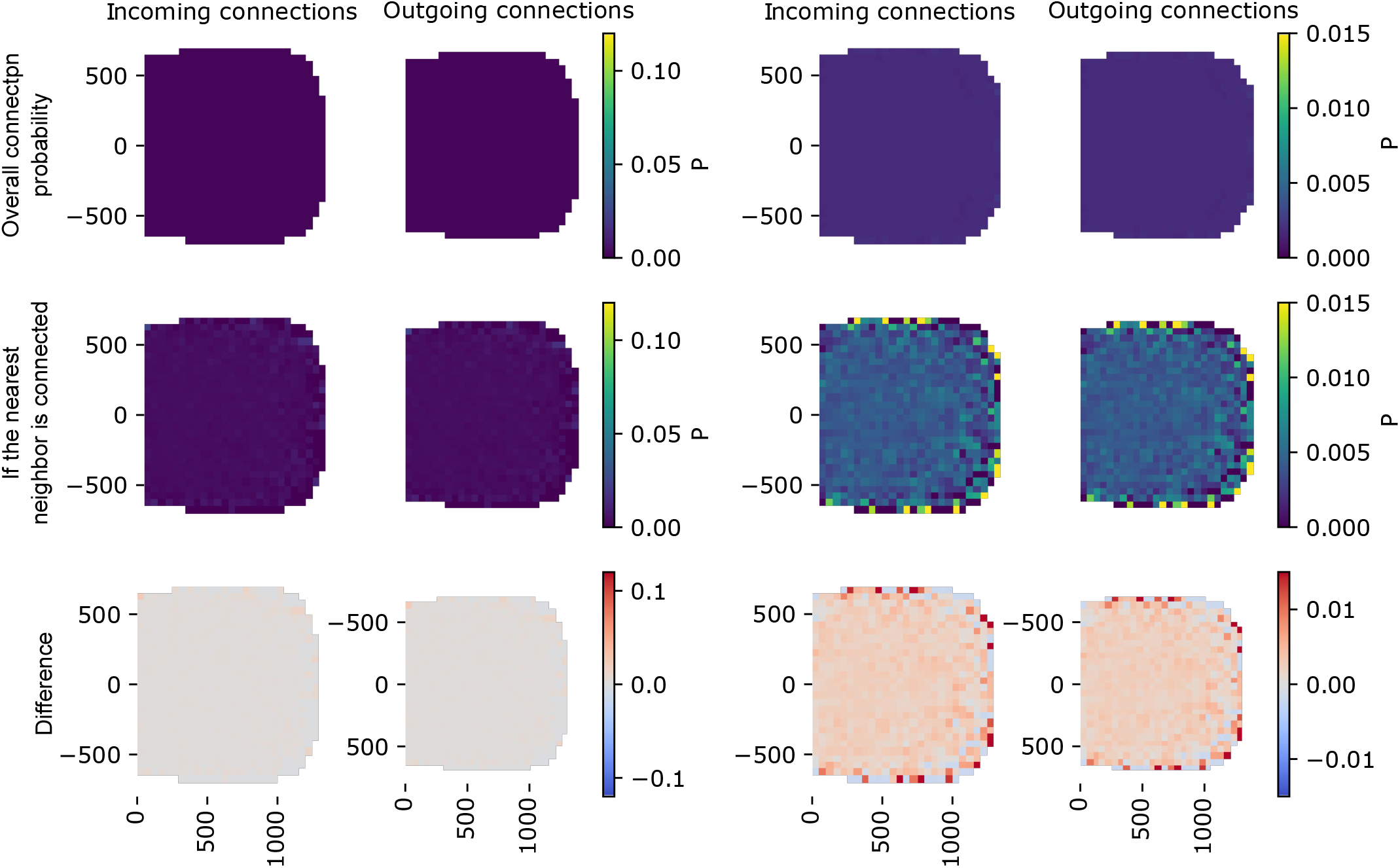
Incoming and outgoing connection probabilities as in Fig. 2 for a preferential attachment model fit to the MICrONS data. Top: Overall connection probability. Middle: Connection probability if the nearest neighbor is connected. Bottom: Difference. Left: Same colormap as in Fig. 2; right: colormap re-scaled.

**Figure S5.**
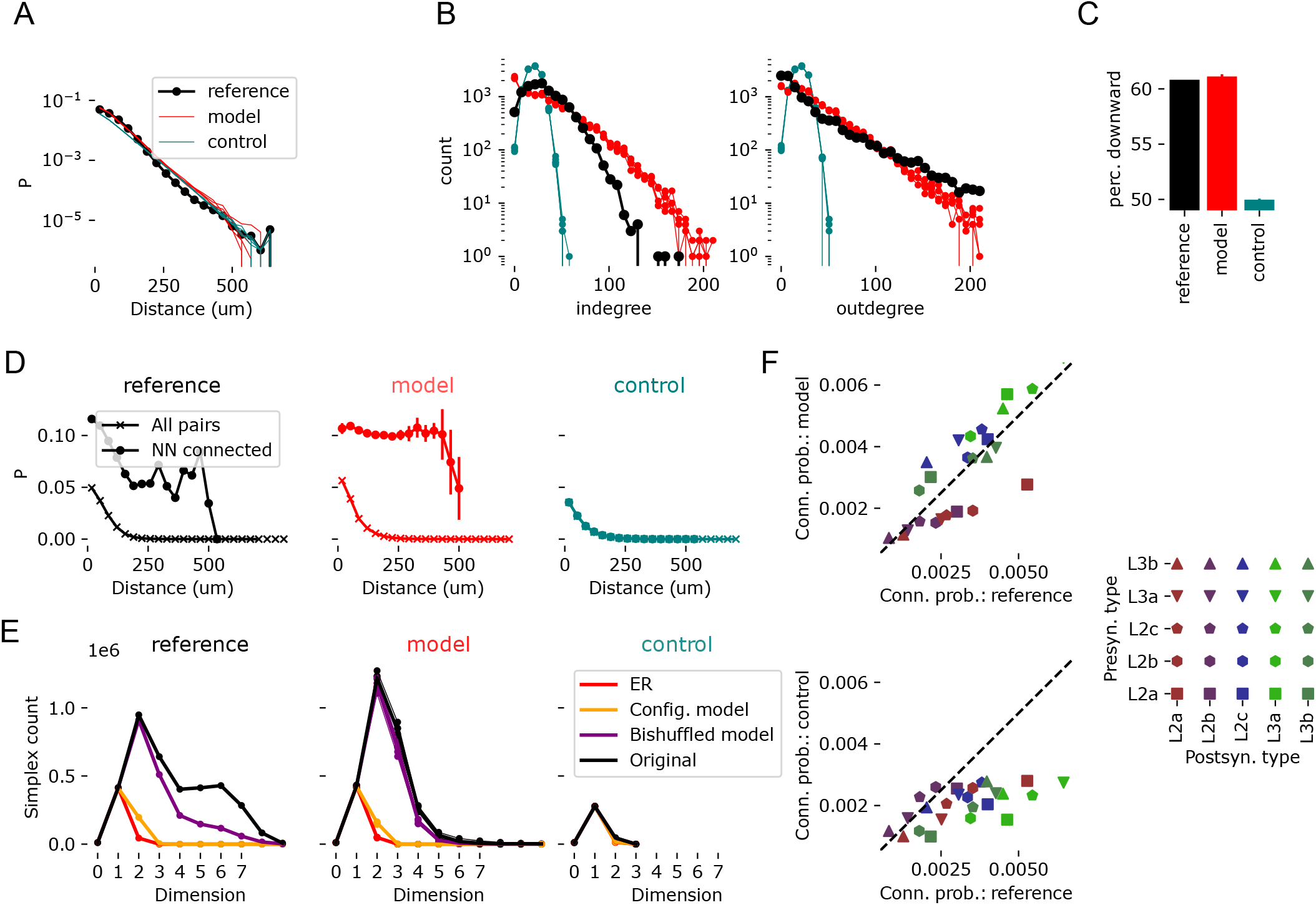
SGSG model compared to excitatory connectivity in layers 2/3 of the MICrONS data. A-E: as in Figure 5, but for layers 2/3 instead of 4/5. F: As in Fig S3. Note that as a test of generalization, parameters *d* and *p* were not re-fitted and *q* was re-fit only to match the total number of edges in this new reference.

**Figure S6.**
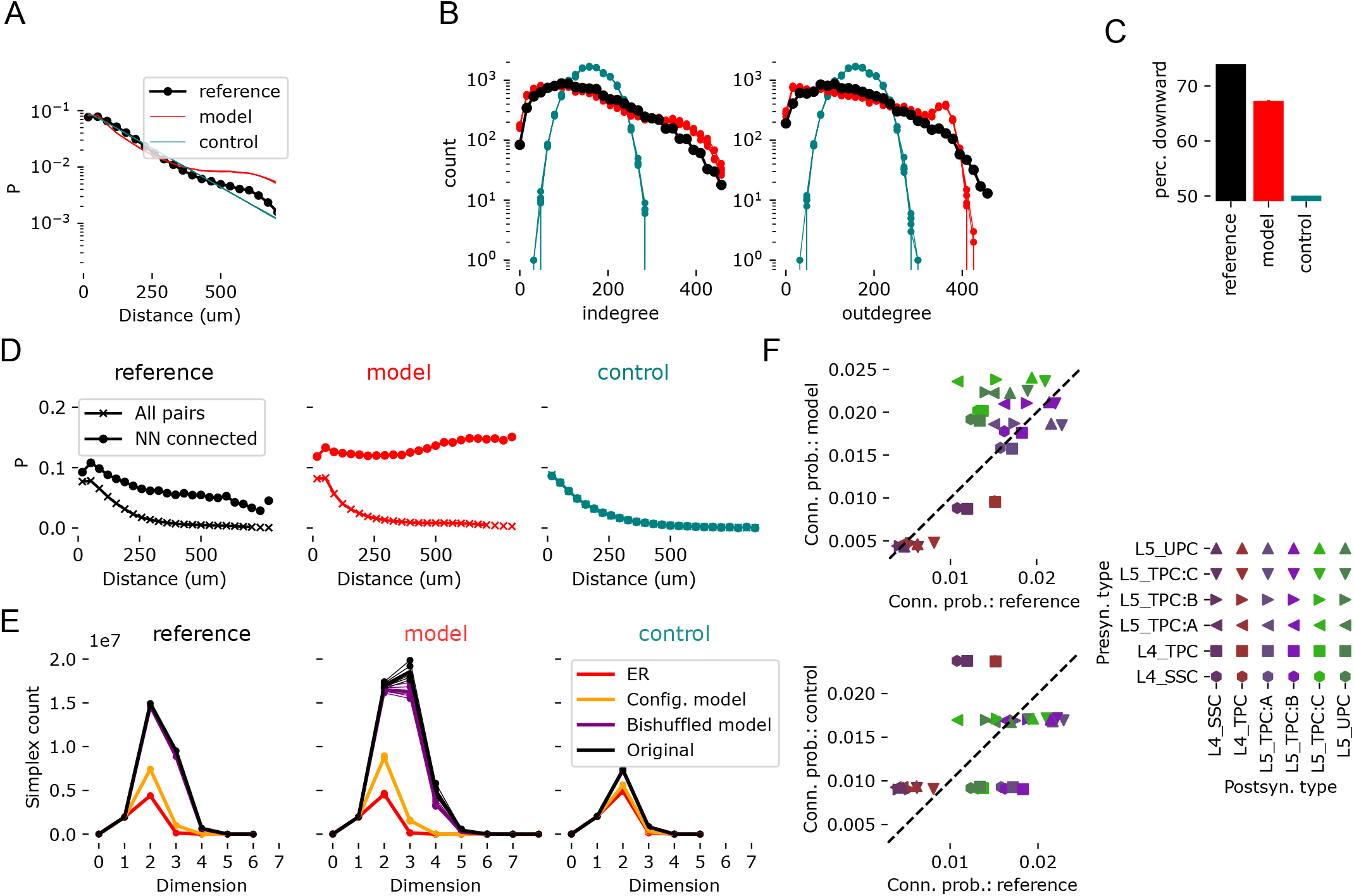
SGSG model compared to excitatory connectivity in layers 4/5 of the rat nbS1 model of Reimann et al. [45] A-E: as in Figure 5, but for the model instead of EM data. F: As in Fig S3.

**Figure S7:**
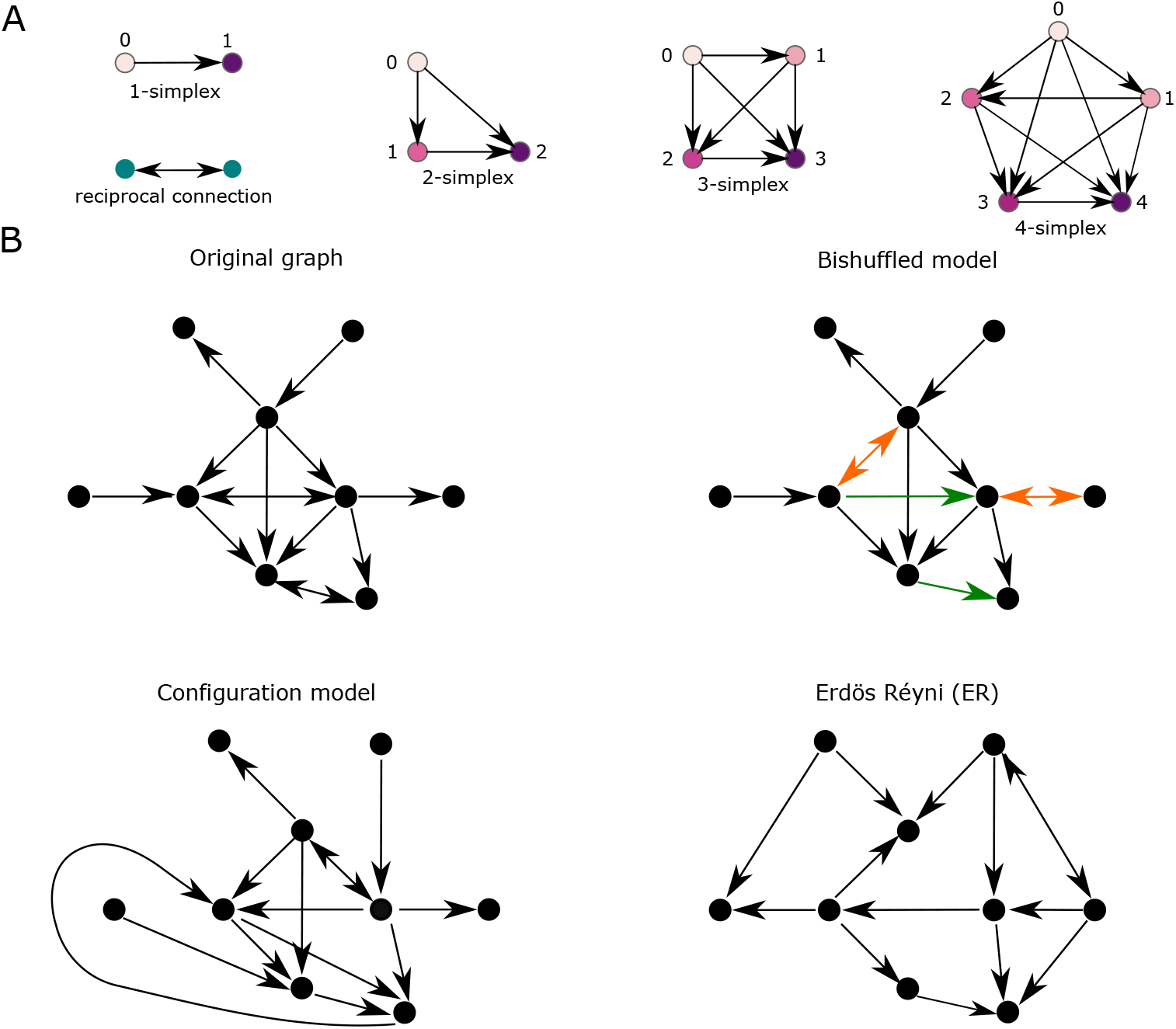
Simplex motifs and stochastic controls. A: Simplex motifs for dimensions 1 through 5. These are fully connected feedforward motifs in which all nodes are connected such that there exists an ordering where there is an edge from *v* to *u* whenever *v < u*. In the diagram, this ordering is both labeled and reflected in the color scheme, transitioning from lighter to darker hues. B: Different stochastic controls of an given original graph (top left). The <u>bishuffled</u> control alters only the positions of reciprocal (bidirectional) connections. Green edges indicate those that were reciprocal in the original, red edges indicate those that become reciprocal in the control. The <u>configuration model</u> and <u>Erdős–Rényi (ER)</u> models generate random graphs with the same number of edges as the original. The configuration model additionally preserves the in-degree and out-degree of each node, while ER scatters them at random across all edges.

**Figure S8.**
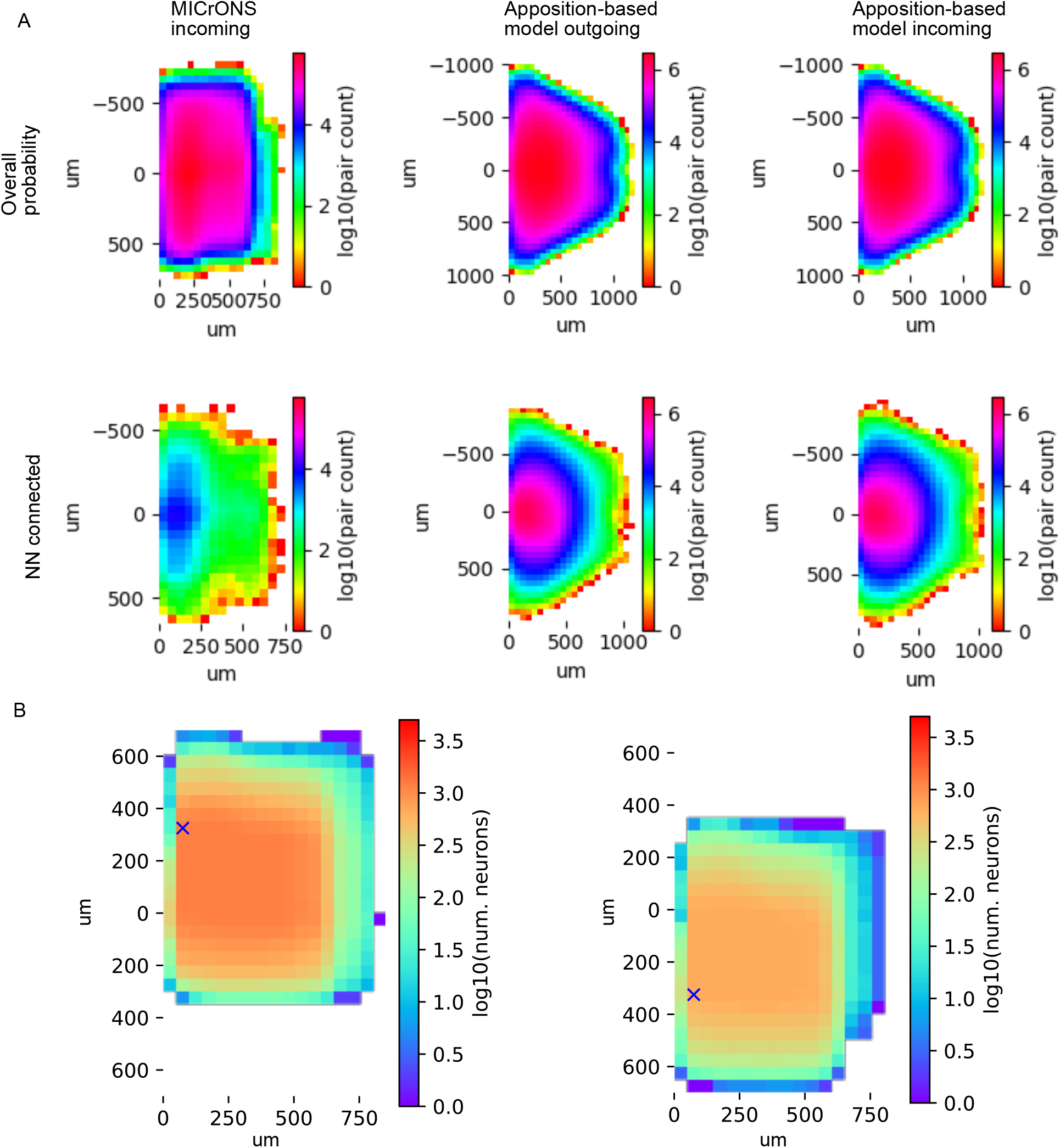
A: Number of pairs sampled for connection probabilities reported in Fig. 3 that are not already reported in the main figure. B: Number of pairs sampled for the connection probability estimates indicated in Fig. 4A, C and the p-values in Fig 4E.

## S2 Supplementary Methods

### S2.1 Offset-dependent connectivity analysis: exploring the effect of the nearest neighbor

#### S2.1.1 Connection probability, conditional on the nearest neighbor connection

Let *M* be the adjacency matrix of a network, such as the MICrONS data. That is, *M*_*u,v*_ is 1 if there is at least one synaptic contact from neuron *u* to neuron *v*, and 0 otherwise. We find for each neuron *u* its *nearest spatial neighbor, n*(*u*) using a KDTree in the implementation of the “scipy” package in python 3.12. We combined *M* with the nearest neighbor information to calculate a matrix *N* such that *N*_*u,v*_ is 1 if there is at least one synaptic contact from neuron *u* to *n*(*v*), and 0 otherwise. The overall connection probability is then the mean value of entries of *M*, excluding entries along the diagonal. The outgoing connection probability conditional on the nearest neighbor connected is the mean value of entries of *M* where the corresponding entry of *N* is 1, excluding diagonal entries and entries where *n*(*v*) = *u*. For the conditional incoming connection probability we simply performed the analysis using the transpose *M*^*T*^ instead. For a distance-dependent analysis we used a matrix of pairwise distances *D* to identify pairs within a given distance bin; for the two-dimensional version we used two matrices *D*_*y*_ and *D*_*x,z*_ denoting the offset along the y-axis and distance in the x,z-plane respectively, as follows.

#### S2.1.2 Connection probability at an offset

We calculated for each pair of neurons their offsets in the horizontal plane and along the vertical axis. Offsets were then binned in both directions with a bin size of 50 × 50 *µm*. Let *B*(*u, v*) be the binning function that associates a pair of nodes with a bin. One can think of a bin as a discretized 2D-vector that describes the offset from the position of *u* to that of *v*. Thus, we refer to a generic bin as 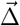.

We computed the outgoing and incoming connection probabilities at a given offset over the entire connectome. That is,

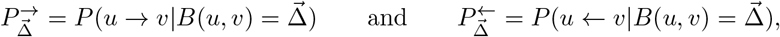

where the arrow represents the presence of an edge in the indicated direction and *u* and *v* run across all nodes in that offset bin. This is a natural extension of the connection probability at a distance to include vertical direction.

Let *NN*(*u*) indicate function that assigns to a node *u* its *nearest neighbor* according to its spatial location.

We calculated the connection probabilities of each bin, conditioned on the nearest neighbor of a neuron being connected in the given offset. That is,

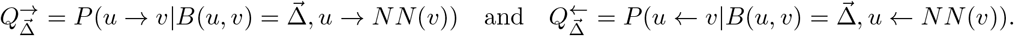

Note in particular, that given our bin size choice, most of the time *u* and *NN*(*u*) belong to the same spatial bin. In this case, 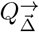 is the probability of more than one connection existing in the offset bin 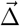, given that one already exists. Therefore, the difference between 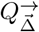 and 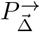 serves as a measure of the degree to which the connectivity in a given bin is statistically non-independent.

#### S2.1.3 Interdependence of connection probability at different offsets

While the above measured statistical dependencies for a given offset, we also measured statistical dependencies between offsets. That is in some sense, to what extent are 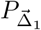 and 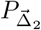 are correlated, for any two of offsets 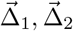.To achieve this, we used a more detailed description of the probability structure, defined at the level of individual neurons.

That is, for a fixed neuron *u* one can calculate its outgoing and incoming connection probabilities at a given offset 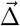

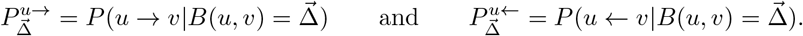

The global connections above, are an average of these values across all neurons *u*. To ensure representative results at the single neuron however, we set the value to be undefined for nodes that had fewer than 100 other nodes at the indicated offset, connected or not.

Let *J*(Δ_1_, Δ_2_) denote the set of nodes *v* for which both 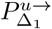 and 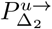 are defined. That is, those nodes that have at least 100 nodes in both offsets from *v* independent of connectivity. Then we can build two vectors

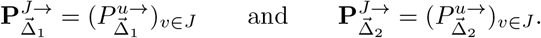

They encode the probability of connectivity for each neuron at each offset. Thus, we calculate their interdependence as the Pearson correlation between them and denote it by 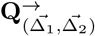. We define 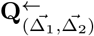 similarly but for incoming connectivity.

To better understand the structure of this measure we clustered the weighted matrices with entries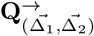 or 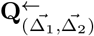 using the Louvain algorithm with a resolution parameter of 1.0. The results were groups of spatial bins that were innervated together by individual neurons (or innervated together individual neurons) more often than expected from the global connection probabilities observed for them.

Additionally, for the plots of Fig. 4C and E, we normalized the connection probability estimates before calculating the correlations. That is, for each neuron we calculated 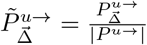, where |*P^u→^*| is the mean outgoing connection probability for that neuron across offsets.

### S2.2 Customizations of SGSGs

#### S2.2.1 Biasing the spatial neighborhood selection

Above, we constructed a simple random geometric graph by placing connections to nodes in the geometric neighborhood of each vertex *v*, i.e., the set of nodes at euclidean distance smaller than *d*, which we denote by *H*_*d*_(*v*), independently at random with probability *p*. We introduce two types of bias into the process. Each bias defines a weight for every edge (*v, w*), which increases or decreases the probability that the pair is selected. These probabilities are then scaled by their corresponding weights and normalized such that the mean probability over all nodes in *H*_*d*_(*n*) remains equal to *p*.

##### Per node biases

The simplest form of bias is to define for each node *v* a tuple of weights (*w*_*o*_(*v*), *w*_*i*_(*v*)), which increase or decrease the relative out- and in-degree of the node *v*. To do so, we bias the probability of choosing each each edge (*v, u*) by the weight *w*_*o*_(*v*) ·*w*_*i*_(*u*). While this bias has potentially different values for each node, in practice we used identical values for large groups of nodes, specifically for nodes representing neurons belonging to the same type.

##### Orientation-based biases

This bias is determined by directional alignment with a base unit vector 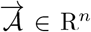, which we call the <u>orientation axis</u>. For two nodes *v* and *u*, corresponding to points **p**_*v*_, **p**_*u*_ ∈ **P**, the probability of forming an edge (*v, u*) is maximally increased when the vector from **p**_*v*_ to **p**_*u*_ is parallel and aligned with 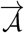 ; it is maximally decreased when the vector points in the opposite direction, and remains unbiased when it is orthogonal to 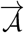.

More precisely, we choose a weight −1≤ *w*_𝒜_ ≤1. Then for any *u H*_*d*_(*v*) let 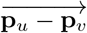 denote the vector from *v* to *u*. Then, the bias weight for (*v, u*) is:

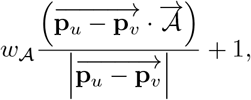

where · denotes the vector dot product. Note that the bias weight ranges from 1 + *w*_𝒜_ to 1 − *w*_𝒜_, depending the level of alignment with 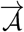.

#### S2.2.2 Reducing stochasticity of *S*_*i*_(*n*)

Above, we describe a process where *S*_*i*+1_(*v*) is generated from *S*_*i*_(*v*) by picking from outgoing neighbors in a random geometric graph independently at random such that the expected size of *S*_*i*+1_(*n*) is *q*. Here, we introduce an alternative method where instead exactly 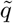 candidates are picked, where 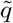 is *q* rounded to the nearest integer. This is similar to the two variants of an Erdos Reyni graph, one with a fixed number of edges and one with a fixed probability of having an edge between a pair of nodes [83; 84]. As there is no variability in the size of *S*_*i*_(*v*), we consider this a less stochastic version of the process. We employ the less stochastic version in the first *k* steps, i.e., for *S*_*i*_ with *i < k*, then switch to the regular version for the remaining steps.

#### S2.2.3 Extension to long-range connectivity

We extend the SGSG to a proof-of-concept version that incorporates additional long-range connections. To achieve this, we generate two random geometric base graphs over the same set of nodes in 3-dimensional space: one with local connectivity constructed as before, and another where distances are defined by transformed coordinates of the original point could, that reflect the structure of long-range connectivity.

More precisely, from a set of points **P** ∈ R^3^, we generate another set of points **P**^∼^∈ R^3^ with transformed *y*-coordinates, while keeping the *x* and *z* coordinates fixed. The modified *y*-coordinates are given by 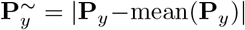, i.e., the distance from an *x, z*-plane at the center of the point cloud.

Then, we build two random geometric graphs, *G*_*d,p*_(**P**) and *G*_*d,p·m*_(**P**^∼^) on the same set of nodes, where *m* is a parameter that determines that balance between the local and long-range edges in the final result. We consider the graph given by the union of edges of *G*_*d,p*_(**P**) and *G*_*d,p·m*_(**P**^∼^) and denote it by 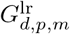(**P,P^∼^**). Finally, we build a stochastic spread graph on it, 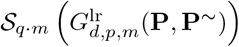 Note that other methods to determine **P**^∼^ and versions in which the point cloud is in *n* dimensional space, are possible and will change the overall and spatial structure of the long-range edges added.

## S3 Characterizing the structure of SGSG with parameter scans

We begin by evaluating this family of graphs with respect to several features of connectivity that characterize the connectivity of cortical networks (Fig. S9). To that end, we built instances of SGSGs while iterating the values of its parameters over a wide range. The parameters were: *d* the maximum distance of connections of the underlying random geometric graph; *p*, the probability that a connection within that distance is added to the random geometric graph; and *q*, the mean number of new nodes that the stochastic process spreads to in each step. Instances were built for 5000 nodes that were randomly uniformly distributed in a cube with a side length of 500*µm*, corresponding to a density of 40,000 per *mm*^3^. Biological cortical cell densities depend on species, but are on the same order of magnitude [85]. We note that the density of the nodes used affects the structure of the SGSG graph in principle, but this can be compensated through the *d* parameter: Note that if we scale each side of the cube and *d* by the same factor *x* and keep the number of points unchanged, we will generate the same SGSGs, but at eight times lower density. Hence, we do not consider the density a parameter to explore.

First, we considered measures of the density of connections: The mean node out-degree as a measure of the overall density of connections; the connection probability within 75*µm* to understand to what degree connections are found specifically between nearby nodes; and the overexpression of reciprocal connectivity (Fig. S9A). Finding e.g., the same mean degree with lower connection probability means that connections are re-distributed to above 75*µm*, indicating different distance-dependence of connectivity. We found that different parameter combinations lead to very different levels of sparsity, indicating that a wide range of types of network can be created. Notably, while the mean degree and probability within 75*µm* were correlated, they were also independent to some degree, therefore showing different types of distance-dependence can be implemented. Significant overexpression of reciprocity was found throughout the parameter space tested, but it was highest in locations of the parameter space resulting in extremely sparse networks. Next, we considered measures of non-random higher-order structure: A common neighbor of a pair of nodes is a third node that is connected to both of them. We define the common neighbor bias (CN bias) as the mean number of common neighbors of connected pairs of nodes divided by the same measure for unconnected pairs of nodes. Its value increases the more the connectivity is clustered. We found that CN bias and in-degree distribution skewness were elevated over values for more random networks throughout the parameter space, although they were again highest for extremely sparse networks. Skewness of out-degrees was relatively constant at values around 2.

To test for the overexpression of motifs in the model, we explored the parameter combination of *d* = 70*µm, p* = 0.1, *q* = 2.5 further, generating 25 SGSG instances. We confirmed distance-dependent, clustered connectivity with log-normal degree distributions (Fig. S9B1-B3). As expected from panel C, the distribution of out-degrees had a longer tail than in-degrees. Further, connected pairs have a larger number of common neighbors at all distances (Fig. 1D3), indicating clustered connectivity. We then focused on the directed simplex motifs that we initially singled out as structurally and functionally relevant in biological neuronal networks and equally present in neuron morphology-derived networks. We found that in the SGSG instances simplices were highly overexpressed compared to strong controls (Fig. S9B4). Notably, their count was higher than even in a bishuffled control (Fig. S7B), that maintains most of the topology of the reference network and only randomizes the locations of bidirectional edges. This indicates that the locations of bidirectional edges are non-random with respect to the simplicial structure and has been previously shown to be the case in biological connectomes [25].

**Figure S9.**
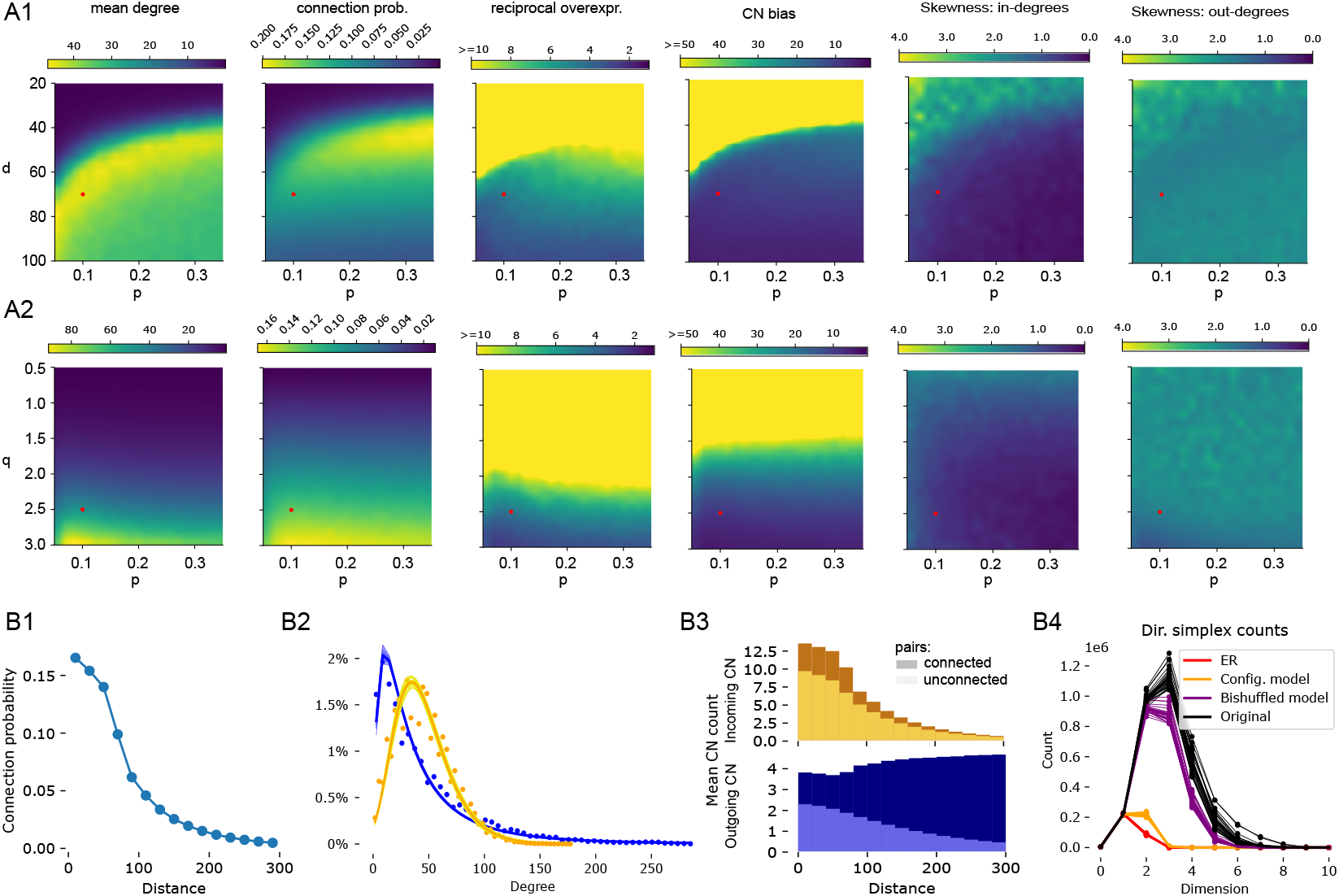
A1: Values of metrics characterizing overall connectivity and higher-order structure of SGSGs with different values of parameters *p* and *d*. For explanation of the parameters and metrics, see Methods. A2: Same, but for parameters *p* and *q*. Red dot indicates location in the parameter space of the instances analyzed further in D. B: More detailed analyses of parameter combination *p* = 0.1, *d* = 70*µm, q* = 2.5. B1: Distance-dependence of connection probability. B2: Distribution of in- (orange) and out- (blue) degrees. Dots indicate mean over 25 instances. Thin lines indicate lognormal fits for individual instances, thick line lognormal fit to pooled data. B3: Common neighbor bias: Number of common graph neighbors in SGSG instances for pairs at indicated distances. Light colored: for unconnected pairs, dark: For connected pairs. Top: incoming, bottom: outgoing common neighbors. Mean over 10 instances indicated. B4: Counts of directed simplex motifs in SGSG instances (black) and various controls fit to it. For details on the controls, see Methods. Each line one instance.

## S4 Comparing other connectome models to the MICrONS connectome

**Figure S10.**
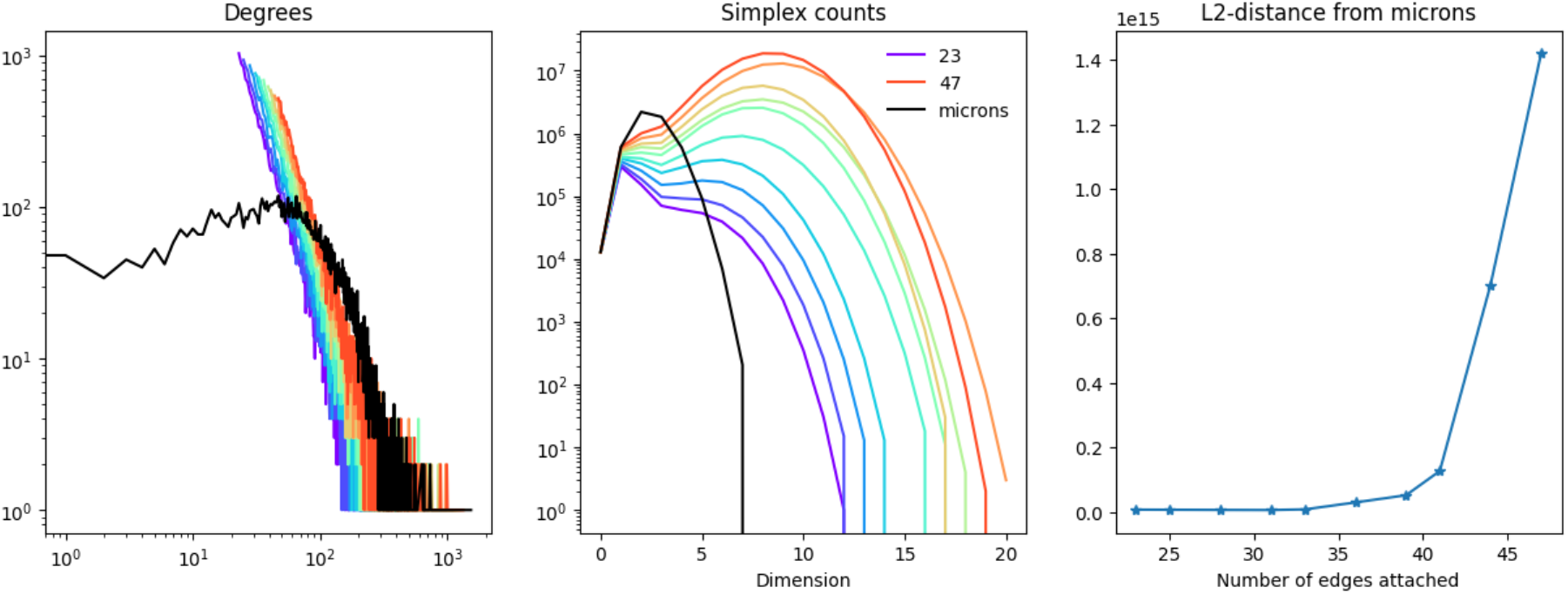
Fitting the undirected model of Barabási and Albert [29] to the MICrONS data. Note that simplex counts in this case are undirected simplices. Different colors indicate different values of the parameter dictating the number of edges (see Methods).

**Figure S11.**
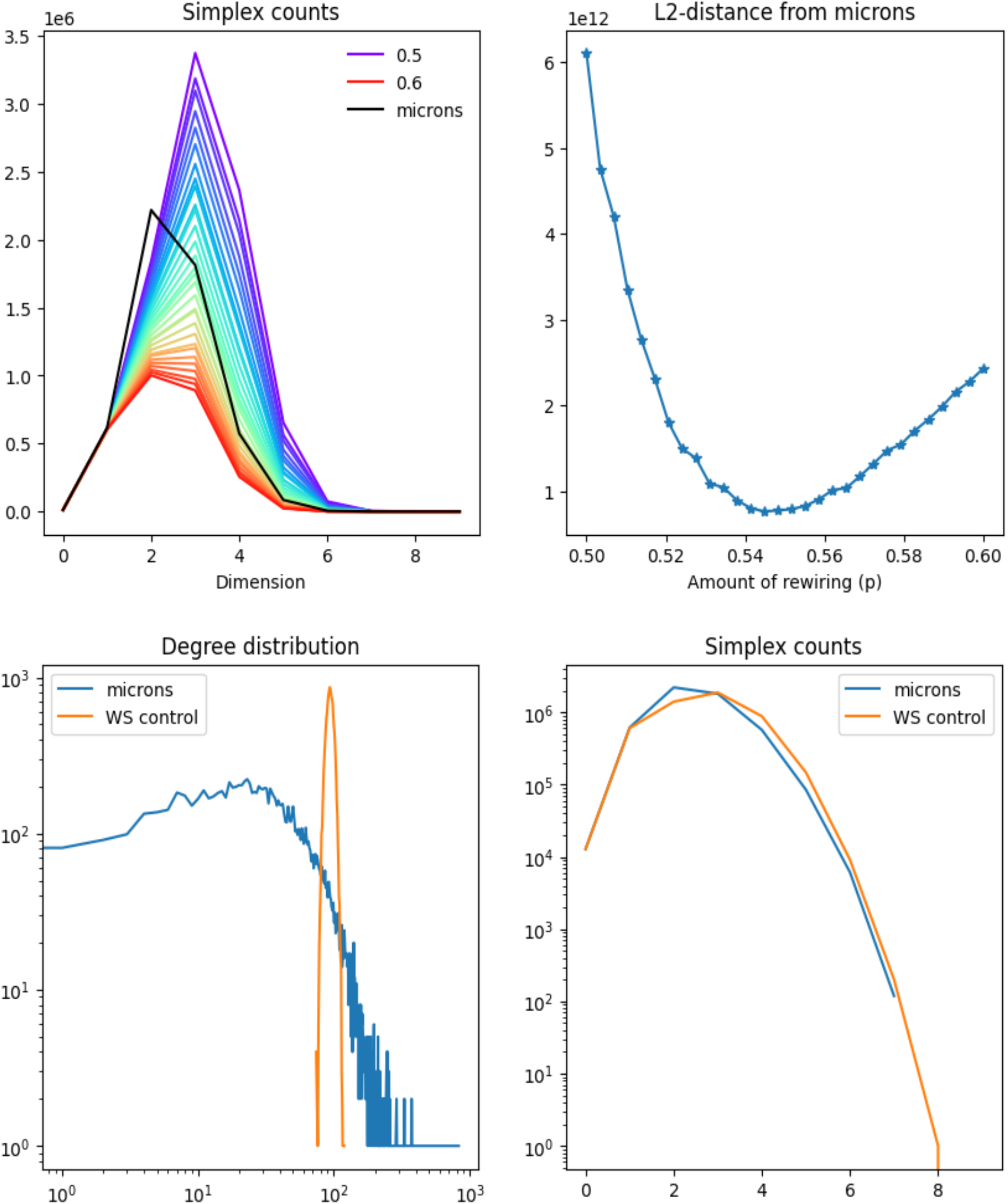
Fitting a scale-free connectome model [32] to the MICrONS data. Top row: Simplex counts and distance of simplex count profile. Bottom right: Simplex counts in logarithmic scale of the best-fitting model. Bottom left: Comparison of degree distributions for the best-fitting model.

**Figure S12.**
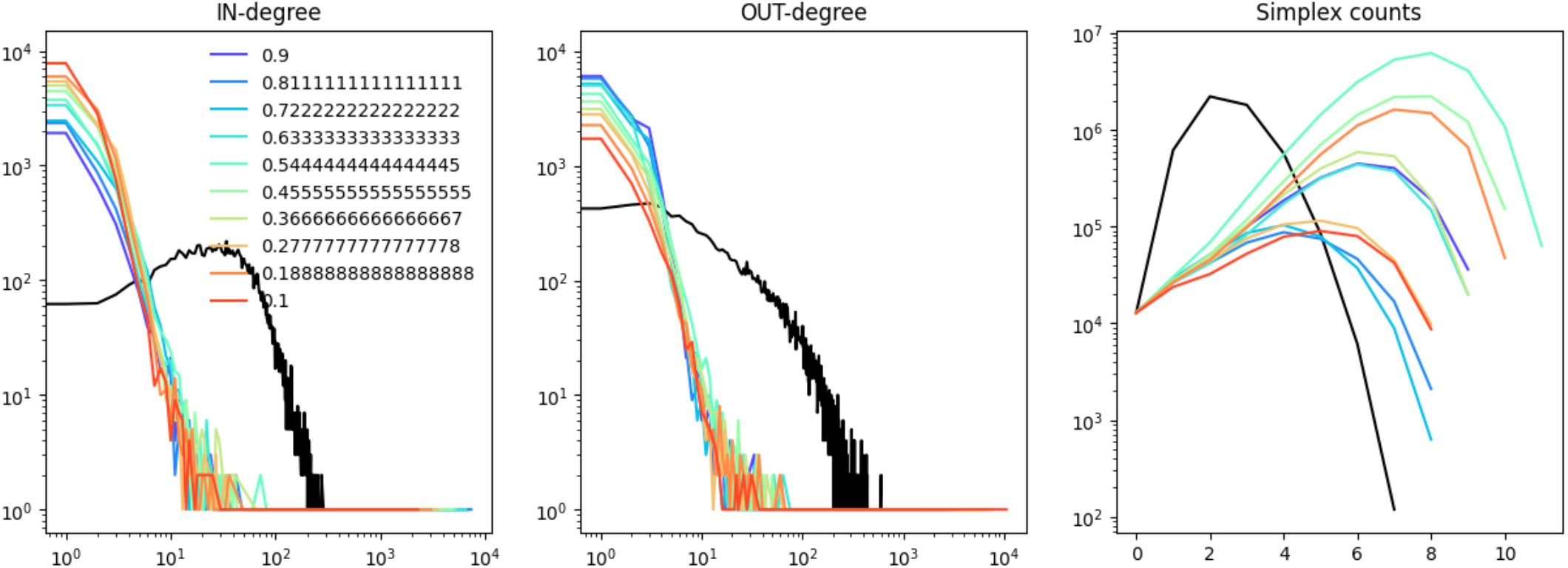
Fitting the directed preferential-attachment (PA) model of Bollobás et al. [82] to the MICrONS data. Left to right: In-degrees, out-degrees and simplex counts. Different colors indicate different paramters for the PA model.

## References

[1] Xinyuan Liang, Lianglong Sun, Xuhong Liao, Tianyuan Lei, Mingrui Xia, Dingna Duan, Zilong Zeng, Qiongling Li, Zhilei Xu, Weiwei Men, et al. Structural connectome architecture shapes the maturation of cortical morphology from childhood to adolescence. Nature Communications, 15 (1):784, 2024.

[2] Jenelle L Wallace and Alex A Pollen. Human neuronal maturation comes of age: cellular mechanisms and species differences. Nature Reviews Neuroscience, 25(1):7–29, 2024.

[3] Elizabeth F Ryder, Lindsey Bullard, Joel Hone, Jonas Olmstead, and Matthew O Ward. Graphical simulation of early development of the cerebral cortex. Computer methods and programs in biomedicine, 59(2):107–114, 1999.

[4] Sarah F Beul, Alexandros Goulas, and Claus C Hilgetag. Comprehensive computational modelling of the development of mammalian cortical connectivity underlying an architectonic type principle. PLoS computational biology, 14(11):e1006550, 2018.

[5] Chenzhong Yin, Xiongye Xiao, Valeriu Balaban, Mikhail E Kandel, Young Jae Lee, Gabriel Popescu, and Paul Bogdan. Network science characteristics of brain-derived neuronal cultures deciphered from quantitative phase imaging data. Scientific reports, 10(1):15078, 2020.

[6] Danke Zhang, Chi Zhang, and Armen Stepanyants. Robust associative learning is sufficient to explain the structural and dynamical properties of local cortical circuits. Journal of Neuroscience, 39(35):6888–6904, 2019.

[7] Yazan N Billeh, Binghuang Cai, Sergey L Gratiy, Kael Dai, Ramakrishnan Iyer, Nathan W Gouwens, Reza Abbasi-Asl, Xiaoxuan Jia, Joshua H Siegle, Shawn R Olsen, et al. Systematic integration of structural and functional data into multi-scale models of mouse primary visual cortex. Neuron, 106(3):388–403, 2020.

[8] James A Roberts, Alistair Perry, Anton R Lord, Gloria Roberts, Philip B Mitchell, Robert E Smith, Fernando Calamante, and Michael Breakspear. The contribution of geometry to the human connectome. Neuroimage, 124:379–393, 2016.

[9] Xiaolong Jiang, Shan Shen, Cathryn R. Cadwell, Philipp Berens, Fabian Sinz, Alexander S. Ecker, Saumil Patel, and Andreas S. Tolias. Principles of connectivity among morphologically defined cell types in adult neocortex. Science, 350(6264):aac9462, November 2015. ISSN 0036-8075, 1095-9203. doi: 10.1126/science.aac9462. URL https://www.science.org/doi/10.1126/science.aac9462.

[10] The MICrONS Consortium, J. Alexander Bae, Mahaly Baptiste, Maya R. Baptiste, Caitlyn A. Bishop, Agnes L. Bodor, Derrick Brittain, Victoria Brooks, JoAnn Buchanan, Daniel J. Bumbarger, Manuel A. Castro, Brendan Celii, Erick Cobos, Forrest Collman, Nuno Maçarico Da Costa, Bethanny Danskin, Sven Dorkenwald, Leila Elabbady, Paul G. Fahey, Tim Fliss, Emmanouil Froudarakis, Jay Gager, Clare Gamlin, William Gray-Roncal, Akhilesh Halageri, James Hebditch, Zhen Jia, Emily Joyce, Justin Ellis-Joyce, Chris Jordan, Daniel Kapner, Nico Kemnitz, Sam Kinn, Lindsey M. Kitchell, Selden Koolman, Kai Kuehner, Kisuk Lee, Kai Li, Ran Lu, Thomas Macrina, Gayathri Mahalingam, Jordan Matelsky, Sarah McReynolds, Elanine Miranda, Eric Mitchell, Shanka Subhra Mondal, Merlin Moore, Shang Mu, Taliah Muhammad, Barak Nehoran, Erika Neace, Oluwaseun Ogedengbe, Christos Papadopoulos, Stelios Papadopoulos, Saumil Patel, Guadalupe Jovita Yasmin Perez Vega, Xaq Pitkow, Sergiy Popovych, Anthony Ramos, R. Clay Reid, Jacob Reimer, Patricia K. Rivlin, Victoria Rose, Zachary M. Sauter, Casey M. Schneider-Mizell, H. Sebastian Seung, Ben Silverman, William Silversmith, Amy Sterling, Fabian H. Sinz, Cameron L. Smith, Rachael Swanstrom, Shelby Suckow, Marc Takeno, Zheng H. Tan, Andreas S. Tolias, Russel Torres, Nicholas L. Turner, Edgar Y. Walker, Tianyu Wang, Adrian Wanner, Brock A. Wester, Grace Williams, Sarah Williams, Kyle Willie, Ryan Willie, William Wong, Jingpeng Wu, Chris Xu, Runzhe Yang, Dimitri Yatsenko, Fei Ye, Wenjing Yin, Rob Young, Szichieh Yu, Daniel Xenes, and Chi Zhang. Functional connectomics spanning multiple areas of mouse visual cortex. Nature, 640(8058):435–447, April 2025. ISSN 0028-0836, 1476-4687. doi: 10.1038/s41586-025-08790-w. URL https://www.nature.com/articles/s41586-025-08790-w.

[11] Ruochen Yang, Frederic Sala, and Paul Bogdan. Hidden network generating rules from partially observed complex networks. Communications Physics, 4(1):199, 2021.

[12] Qing Huang, Yijun Chen, Shijie Liu, Cheng Xu, Tingting Cao, Yongchao Xu, Xiaojun Wang, Gong Rao, Anan Li, Shaoqun Zeng, et al. Weakly supervised learning of 3d deep network for neuron reconstruction. Frontiers in Neuroanatomy, 14:38, 2020.

[13] Sven Dorkenwald, Claire E McKellar, Thomas Macrina, Nico Kemnitz, Kisuk Lee, Ran Lu, Jingpeng Wu, Sergiy Popovych, Eric Mitchell, Barak Nehoran, et al. Flywire: online community for whole-brain connectomics. Nature methods, 19(1):119–128, 2022.

[14] Cassandra Hoffmann, Ellie Cho, Andrew Zalesky, and Maria A Di Biase. From pixels to connections: exploring in vitro neuron reconstruction software for network graph generation. Communications Biology, 7(1):571, 2024.

[15] Brendan Celii, Stelios Papadopoulos, Zhuokun Ding, Paul G Fahey, Eric Wang, Christos Papadopoulos, Alexander B Kunin, Saumil Patel, J Alexander Bae, Agnes L Bodor, et al. Neurd offers automated proofreading and feature extraction for connectomics. Nature, 640(8058):487–496, 2025.

[16] Sven Dorkenwald, Casey M Schneider-Mizell, Derrick Brittain, Akhilesh Halageri, Chris Jordan, Nico Kemnitz, Manual A Castro, William Silversmith, Jeremy Maitin-Shephard, Jakob Troidl, et al. Cave: Connectome annotation versioning engine. Nature Methods, pages 1–9, 2025.

[17] Sen Song, Per Jesper Sjöström, Markus Reigl, Sacha Nelson, and Dmitri B Chklovskii. Highly Nonrandom Features of Synaptic Connectivity in Local Cortical Circuits. PLoS Biology, 3(3):e68, March 2005. ISSN 1545-7885. doi: 10.1371/journal.pbio.0030068. URL https://dx.plos.org/10.1371/journal.pbio.0030068.

[18] R. Perin, T. K. Berger, and H. Markram. A synaptic organizing principle for cortical neuronal groups. Proceedings of the National Academy of Sciences, 108(13):5419–5424, March 2011. ISSN 0027-8424, 1091-6490. doi: 10.1073/pnas.1016051108. URL http://www.pnas.org/cgi/doi/10.1073/pnas.1016051108.

[19] Emma K Towlson, Petra E Vértes, Sebastian E Ahnert, William R Schafer, and Edward T Bullmore. The rich club of the c. elegans neuronal connectome. J. Neurosci., 33(15):6380–6387, 2013. doi: 10.1523/JNEUROSCI.3784-12.2013.

[20] Ann E Sizemore, Chad Giusti, Ari Kahn, Jean M Vettel, Richard F Betzel, and Danielle S Bassett. Cliques and cavities in the human connectome. J. Comput. Neurosci., 44:115–145, 2018. doi: 10.1007/s10827-017-0672-6.

[21] Bosiljka Tadić, Miroslav Andjelković, and Roderick Melnik. Functional geometry of human connectomes. Sci. Rep., 9(1):12060, 2019. doi: 10.1038/s41598-019-48568-5.

[22] Ann E Sizemore, Jennifer E Phillips-Cremins, Robert Ghrist, and Danielle S Bassett. The importance of the whole: topological data analysis for the network neuroscientist. Netw. Neurosci., 3 (3):656–673, 2019. doi: 10.1162/netna00073.

[23] Miroslav Andjelković, Bosiljka Tadić, and Roderick Melnik. The topology of higher-order complexes associated with brain hubs in human connectomes. Sci. Rep., 10(1):17320, 2020. doi: 10.1038/s41598-020-74392-3.

[24] Dinghua Shi, Zhifeng Chen, Xiang Sun, Qinghua Chen, Chuang Ma, Yang Lou, and Guanrong Chen. Computing cliques and cavities in networks. Commun. Phys., 4(1):249, 2021. doi: 10.1038/s42005-021-00748-4.

[25] Daniela Egas Santander, Christoph Pokorny, András Ecker, Jānis Lazovskis, Matteo Santoro, Jason P. Smith, Kathryn Hess, Ran Levi, and Michael W. Reimann. Heterogeneous and higher-order cortical connectivity undergirds efficient, robust, and reliable neural codes. iScience, 28(1):111585, January 2025. ISSN 25890042. doi: 10.1016/j.isci.2024.111585. URL https://linkinghub.elsevier.com/retrieve/pii/S2589004224028128.

[26] Y. Peng, A. Bjelde, P.V. Aceituno, F.X. Mittermaier, H. Planert, S. Grosser, J. Onken, K. Faust, T. Kalbhenn, M. Simon, H. Radbruch, P. Fidzinski, D. Schmitz, H. Alle, M. Holtkamp, I. Vida, B.F. Grewe, and J.R.P. Geiger. Directed and acyclic synaptic connectivity in the human layer 2-3 cortical microcircuit. Science, 2024. doi: 10.1126/science.adg8828.

[27] Federico Tesler, Roberta Maria Lorenzi, Adam Ponzi, Claudia Casellato, Fulvia Palesi, Daniela Gandolfi, Claudia AM Gandini Wheeler Kingshott, Jonathan Mapelli, Egidio d’Angelo, Michele Migliore, et al. Multiscale modeling of neuronal dynamics in hippocampus ca1. Frontiers in Computational Neuroscience, 18:1432593, 2024.

[28] Daniela Gandolfi, Jonathan Mapelli, Sergio Solinas, Robin De Schepper, Alice Geminiani, Claudia Casellato, Egidio D’Angelo, and Michele Migliore. A realistic morpho-anatomical connection strategy for modelling full-scale point-neuron microcircuits. Scientific Reports, 12(1):13864, 2022.

[29] Albert-László Barabási and Réka Albert. Emergence of scaling in random networks. science, 286 (5439):509–512, 1999.

[30] Tom Britton. Directed preferential attachment models. arXiv preprint arXiv:1810.02715, 2018.

[31] Nicolas Brunel. Is cortical connectivity optimized for storing information? Nature neuroscience, 19(5):749–755, 2016.

[32] Duncan J Watts and Steven H Strogatz. Collective dynamics of ‘small-world’networks. nature, 393(6684):440–442, 1998.

[33] Abolfazl Ramezanpour and V Karimipour. Simple models of small-world networks with directed links. Physical review E, 66(3):036128, 2002.

[34] Miguel Ángel García-Cabezas and Basilis Zikopoulos. Evolution, development, and organization of the cortical connectome. PLoS biology, 17(5):e3000259, 2019.

[35] Anastasiya Salova and István A Kovács. Combined topological and spatial constraints are required to capture the structure of neural connectomes. Network Neuroscience, 9(1):181–206, 2025.

[36] Danyal Akarca, Alexander WE Dunn, Philipp J Hornauer, Silvia Ronchi, Michele Fiscella, Congwei Wang, Marco Terrigno, Ravi Jagasia, Petra E Vértes, Susanna B Mierau, Ole Paulsen, Stephen J Eglen, Andreas Hierlemann, Duncan E Astle, and Manuel Schröter. Homophilic wiring principles underpin neuronal network topology in vitro. eLife, 14:e85300, jul 2025. ISSN 2050-084X. doi: 10.7554/eLife.85300. URL https://doi.org/10.7554/eLife.85300.

[37] Ruochen Yang and Paul Bogdan. Controlling the multifractal generating measures of complex networks. Scientific reports, 10(1):5541, 2020.

[38] Xiongye Xiao, Hanlong Chen, and Paul Bogdan. Deciphering the generating rules and functionalities of complex networks. Scientific reports, 11(1):22964, 2021.

[39] Xinhua Zhan, Glen C C. Jickling, Bradley P. Ander, Dazhi Liu, Boryana Stamova, Christopher Cox, Lee-Way Jin, Charles DeCarli, and Frank R. Sharp. Myelin injury and degraded myelin vesicles in alzheimer’s disease. Current Alzheimer Research, 11(3):232–238, 2014.

[40] J Modolo, M Hassan, F Wendling, and P Benquet. Decoding the circuitry of consciousness: from local microcircuits to brain-scale networks. network neuroscience 4 (2): 315–337, 2020.

[41] András Ecker, Daniela Egas Santander, Marwan Abdellah, Jorge Blanco Alonso, Sirio Bolaños-Puchet, Giuseppe Chindemi, Dhuruva Priyan Gowri Mariyappan, James B Isbister, James Gonzalo King, Pramod Kumbhar, Ioannis Magkanaris, Eilif B Muller, and Michael W Reimann. Assemblies, synapse clustering and network topology interact with plasticity to explain structure-function relationships of the cortical connectome, November 2024. URL https://elifesciences.org/reviewed-preprints/101850v1.

[42] William A Catterall, Indira M Raman, Hugh PC Robinson, Terrence J Sejnowski, and Ole Paulsen. The hodgkin-huxley heritage: from channels to circuits. Journal of Neuroscience, 32(41):14064–14073, 2012.

[43] Armen Stepanyants and Dmitri B Chklovskii. Neurogeometry and potential synaptic connectivity. Trends in neurosciences, 28(7):387–394, 2005.

[44] Tom Binzegger, Rodney J Douglas, and Kevan AC Martin. A quantitative map of the circuit of cat primary visual cortex. Journal of Neuroscience, 24(39):8441–8453, 2004.

[45] Michael W Reimann, Sirio Bolaños-Puchet, Jean-Denis Courcol, Daniela Egas Santander, Alexis Arnaudon, Benoît Coste, Fabien Delalondre, Thomas Delemontex, Adrien Devresse, Hugo Dictus, Alexander Dietz, András Ecker, Cyrille Favreau, Gianluca Ficarelli, Mike Gevaert, Joni Herttuainen, James B Isbister, Lida Kanari, Daniel Keller, James King, Pramod Kumbhar, Samuel Lapere, Jānis Lazovskis, Huanxiang Lu, Nicolas Ninin, Fernando Pereira, Judit Planas, Christoph Pokorny, Juan Luis Riquelme, Armando Romani, Ying Shi, Jason P Smith, Vishal Sood, Mohit Srivastava, Werner Van Geit, Liesbeth Vanherpe, Matthias Wolf, Ran Levi, Kathryn Hess, Felix Schürmann, Eilif B Muller, Henry Markram, and Srikanth Ramaswamy. Modeling and Simulation of Neocortical Micro-and Mesocircuitry. Part I: Anatomy, August 2024. URL https://elifesciences.org/reviewed-preprints/99688v1.

[46] Marcel Oberlaender, Christiaan PJ De Kock, Randy M Bruno, Alejandro Ramirez, Hanno S Meyer, Vincent J Dercksen, Moritz Helmstaedter, and Bert Sakmann. Cell type–specific three-dimensional structure of thalamocortical circuits in a column of rat vibrissal cortex. Cerebral cortex, 22(10):2375–2391, 2012.

[47] Daniel Udvary, Philipp Harth, Jakob H Macke, Hans-Christian Hege, Christiaan PJ de Kock, Bert Sakmann, and Marcel Oberlaender. The impact of neuron morphology on cortical network architecture. Cell Reports, 39(2), 2022.

[48] Christopher L Rees, Keivan Moradi, and Giorgio A Ascoli. Weighing the evidence in peters’ rule: does neuronal morphology predict connectivity? Trends in neurosciences, 40(2):63–71, 2017.

[49] Eyal Gal, Michael London, Amir Globerson, Srikanth Ramaswamy, Michael W Reimann, Eilif Muller, Henry Markram, and Idan Segev. Rich cell-type-specific network topology in neocortical microcircuitry. Nature neuroscience, 20(7):1004–1013, 2017.

[50] C.M. Schneider-Mizell, A.L. Bodor, D. Brittain, J. Buchanan, D.J. Bumbarger, L. Elabbady, C. Gamlin, D. Kapner, S. Kinn, G. Mahalingam, S. Seshamani, S. Suckow, M. Takeno, R. Torres, W. Yin, S. Dorkenwald, J.A. Bae, M.A. Castro, A. Halageri, Z. Jia, C. Jordan, N. Kemnitz, K. Lee, K. Li, R. Lu, T. Macrina, E. Mitchell, S.S. Mondal, S. Mu, B. Nehoran, S. Popovych, W. Silversmith, N.L. Turner, W. Wong, J. Wu, MICrONS Consortium, J. Reimer, A.S. Tolias, H.S. Seung, R.C. Reid, F. Collman, and N. Maçarico da Costa. Cell-type-specific inhibitory circuitry from a connectomic census of mouse visual cortex. bioRxiv [Preprint], 2024. doi: 10.1101/2023.01.23.525290.

[51] Leila Elabbady, Sharmishtaa Seshamani, Shang Mu, Gayathri Mahalingam, Casey M Schneider-Mizell, Agnes L Bodor, J Alexander Bae, Derrick Brittain, JoAnn Buchanan, Daniel J Bumbarger, et al. Perisomatic ultrastructure efficiently classifies cells in mouse cortex. Nature, 640(8058): 478–486, 2025.

[52] Ben Piazza, Dániel L. Barabási, André Ferreira Castro, Giulia Menichetti, and Albert-László Barabási. Physical Network Constraints Define the Lognormal Architecture of the Brain’s Connectome, February 2025. URL http://biorxiv.org/lookup/doi/10.1101/2025.02.27.640551.

[53] Michael J Barber. Modularity and community detection in bipartite networks. Physical Review E—Statistical, Nonlinear, and Soft Matter Physics, 76(6):066102, 2007.

[54] Vernon B Mountcastle. The columnar organization of the neocortex. Brain: a journal of neurology, 120(4):701–722, 1997.

[55] Jean-Gabriel Young, Giovanni Petri, Francesco Vaccarino, and Alice Patania. Construction of and efficient sampling from the simplicial configuration model. Phys. Rev. E, 96(3):032312, 2017. doi: 10.1103/PhysRevE.96.032312.

[56] Florian Unger and Jonathan Krebs. Mcmc sampling of directed flag complexes with fixed undirected graphs. Journal of Applied and Computational Topology, 8(6):1881–1916, 2024.

[57] Armen Stepanyants, Luis M Martinez, Alex S Ferecskó, and Zoltán F Kisvárday. The fractions of short-and long-range connections in the visual cortex. Proceedings of the National Academy of Sciences, 106(9):3555–3560, 2009.

[58] Yuanzhe Liu, Caio Seguin, Richard F Betzel, Daniel Han, Danyal Akarca, Maria A Di Biase, and Andrew Zalesky. A generative model of the connectome with dynamic axon growth. Network Neuroscience, 8(4):1192–1211, 2024.

[59] Frederic Zubler and Rodney Douglas. A framework for modeling the growth and development of neurons and networks. Frontiers in computational neuroscience, 3:757, 2009.

[60] Eleni Vasilaki and Michele Giugliano. Emergence of connectivity motifs in networks of model neurons with short-and long-term plastic synapses. PloS one, 9(1):e84626, 2014.

[61] Michael W Reimann, Daniela Egas Santander, András Ecker, and Eilif B Muller. Specific inhibition and disinhibition in the higher-order structure of a cortical connectome. Cerebral Cortex, 34 (11):bhae433, November 2024. ISSN 1047-3211, 1460-2199. doi: 10.1093/cercor/bhae433. URL https://academic.oup.com/cercor/article/doi/10.1093/cercor/bhae433/7889117.

[62] K Ruby, K Falvey, and RJ Kulesza. Abnormal neuronal morphology and neurochemistry in the auditory brainstem of fmr1 knockout rats. Neuroscience, 303:285–298, 2015.

[63] Yan-Bin Shi, Tian Tu, Juan Jiang, Qi-Lei Zhang, Jia-Qi Ai, Aihua Pan, Jim Manavis, Ewen Tu, and Xiao-Xin Yan. Early dendritic dystrophy in human brains with primary age-related tauopathy. Frontiers in Aging Neuroscience, 12:596894, 2020.

[64] Robert F Berman, Karl D Murray, Gloria Arque, Michael R Hunsaker, and H Jürgen Wenzel. Abnormal dendrite and spine morphology in primary visual cortex in the cgg knock-in mouse model of the fragile x premutation. Epilepsia, 53:150–160, 2012.

[65] Gaynor Smith, Sean T Sweeney, Cahir J O’Kane, and Andreas Prokop. How neurons maintain their axons long-term: an integrated view of axon biology and pathology. Frontiers in Neuroscience, 17:1236815, 2023.

[66] Ruochen Yang, Heng Ping, Xiongye Xiao, Roozbeh Kiani, and Paul Bogdan. Spiking dynamics of individual neurons reflect changes in the structure and function of neuronal networks. Nature Communications, 16(1):6994, 2025.

[67] Ruochen Yang, Gaurav Gupta, and Paul Bogdan. Data-driven perception of neuron point process with unknown unknowns. In Proceedings of the 10th ACM/IEEE International Conference on Cyber-Physical Systems, pages 259–269, 2019.

[68] Mingdong Li, Shuhang Chen, Xiang Zhang, and Yiwen Wang. Neural correlation integrated adaptive point process filtering on population spike trains. IEEE Transactions on Neural Systems and Rehabilitation Engineering, 2025.

[69] Gaurav Gupta, Justin Rhodes, Roozbeh Kiani, and Paul Bogdan. Neuron particles capture network topology and behavior from single units. bioRxiv, pages 2021–12, 2021.

[70] Tommaso Boccato, Matteo Ferrante, Andrea Duggento, and Nicola Toschi. Beyond multilayer perceptrons: Investigating complex topologies in neural networks. Neural Networks, 171:215–228, 2024.

[71] Sofia Leite, Bruno Mota, António Ramos Silva, Michael Lamport Commons, Patrice Marie Miller, and Pedro Pereira Rodrigues. Hierarchical growth in neural networks structure: Organizing inputs by order of hierarchical complexity. Plos one, 18(8):e0290743, 2023.

[72] Bosiljka Tadić and Roderick Melnik. Self-organised critical dynamics as a key to fundamental features of complexity in physical, biological, and social networks. Dynamics, 1(2):181–197, 2021. doi: 10.3390/dynamics1020011.

[73] Bengier Ü lgen Kilic and Dane Taylor. Simplicial cascades are orchestrated by the multidimensional geometry of neuronal complexes. Commun. Phys., 5(1):278, 2022. doi: 10.1038/s42005-022-01062-3.

[74] Marc Montalà-Flaquer, Clara F López-León, Daniel Tornero, Akke Mats Houben, Tanguy Fardet, Pascal Monceau, Samuel Bottani, and Jordi Soriano. Rich dynamics and functional organization on topographically designed neuronal networks in vitro. Iscience, 25(12), 2022.

[75] Akke Mats Houben, Jordi Garcia-Ojalvo, and Jordi Soriano. Role of connectivity anisotropies in the dynamics of cultured neuronal networks. PLOS Computational Biology, 21(11):e1012727, 2025.

[76] Dmitri B Chklovskii, Thomas Schikorski, and Charles F Stevens. Wiring optimization in cortical circuits. Neuron, 34(3):341–347, 2002.

[77] Nicholas J Priebe. Mechanisms of orientation selectivity in the primary visual cortex. Annual review of vision science, 2(1):85–107, 2016.

[78] Zizhen Yao, Cindy TJ Van Velthoven, Michael Kunst, Meng Zhang, Delissa McMillen, Changkyu Lee, Won Jung, Jeff Goldy, Aliya Abdelhak, Matthew Aitken, et al. A high-resolution transcriptomic and spatial atlas of cell types in the whole mouse brain. Nature, 624(7991):317–332, 2023.

[79] Seung Wook Oh, Julie A. Harris, Lydia Ng, Brent Winslow, Nicholas Cain, Stefan Mihalas, Quanxin Wang, Chris Lau, Leonard Kuan, Alex M. Henry, Marty T. Mortrud, Benjamin Ouellette, Thuc Nghi Nguyen, Staci A. Sorensen, Clifford R. Slaughterbeck, Wayne Wakeman, Yang Li, David Feng, Anh Ho, Eric Nicholas, Karla E. Hirokawa, Phillip Bohn, Kevin M. Joines, Hanchuan Peng, Michael J. Hawrylycz, John W. Phillips, John G. Hohmann, Paul Wohnoutka, Charles R. Gerfen, Christof Koch, Amy Bernard, Chinh Dang, Allan R. Jones, and Hongkui Zeng. A mesoscale connectome of the mouse brain. Nature, 508(7495):207–214, April 2014. ISSN 0028-0836, 1476-4687. doi: 10.1038/nature13186. URL https://www.nature.com/articles/nature13186.

[80] Gidon Levakov, Joshua Faskowitz, Galia Avidan, and Olaf Sporns. Mapping individual differences across brain network structure to function and behavior with connectome embedding. NeuroImage, 242:118469, 2021.

[81] Omer Bobrowski and Matthew Kahle. Topology of random geometric complexes: a survey. Journal of applied and Computational Topology, 1:331–364, 2018.

[82] Béla Bollobás, Christian Borgs, Jennifer T Chayes, and Oliver Riordan. Directed scale-free graphs. In SODA, volume 3, pages 132–139. Baltimore, MD, United States, 2003.

[83] Erdos Paul. On random graphs. Publicationes Mathematicaes, 6:290–297, 1959.

[84] Edgar N Gilbert. Random graphs. The Annals of Mathematical Statistics, 30(4):1141–1144, 1959.

[85] Diana Furcila, Marta Domínguez-Álvaro, Javier DeFelipe, and Lidia Alonso-Nanclares. Subregional Density of Neurons, Neurofibrillary Tangles and Amyloid Plaques in the Hippocampus of Patients With Alzheimer’s Disease. Frontiers in Neuroanatomy, 13:99, December 2019. ISSN 1662-5129. doi: 10.3389/fnana.2019.00099. URL https://www.frontiersin.org/article/10.3389/fnana.2019.00099/full.

